# Are protein-ligand complexes robust structures?

**DOI:** 10.1101/454165

**Authors:** Maciej Majewski, Sergio Ruiz-Carmona, Xavier Barril

## Abstract

The predominant view in structure-based drug design is that small-molecule ligands, once bound to their target structures, display a well-defined binding mode. While this is convenient from a design perspective, it ignores the fact that structural stability (robustness) is not necessary for thermodynamic stability (binding affinity). In fact, any potential benefit of a rigid binding mode will have to be balanced against the entropic penalty that it entails. Surprisingly, little is known about the causes, consequences and real degree of robustness of protein-ligand complexes. Here we investigate two diverse sets of structures, comprising 79 drug-like and 27 fragment ligands, respectively. We focus on hydrogen bond interactions (469 in total), as they have been described as essential for structural stability. We find that 75% of complexes are anchored by at least one robust hydrogen bond, the remaining 25% either form loose complexes or are constrained by other interactions types. The first type of complexes generally combine a single anchoring point with looser regions, thus balancing order and disorder. Completely constricted protein-ligand complexes are rare and seem to fulfil a functional necessity. Structural stability analysis reveals a hidden layer of complexity in protein-ligand complexes that should be considered in ligand design.

## INTRODUCTION

Biomolecular systems present a large number of degrees of freedom and must find a suitable balance between order and disorder. In the particular case of non-covalent complexes, they can exist in a continuum spectrum of possibilities, ranging from the lock-and-key model to extreme disorder.^1,2^ While the importance of target flexibility is well-appreciated in drug discovery,^3^ the flexibility of small-molecule ligands in their bound state has attracted much less attention. Detailed analyses reveal that ligands often retain residual mobility.^4–6^ However, changes in binding mode are more the exception than the norm^7,8^ and ligand design based on rigid crystallographic geometries has been remarkably successful.^9^ Perhaps for this reason, little is known about the molecular mechanisms that control structural stability, to what extent do ligands preserve flexibility or what are the energetic and functional consequences of rigidity.

It is important to note that structural stability (robustness) is fundamentally different from thermodynamic stability (i.e. binding free energy; *ΔG*_*bind*_). This is eloquently exemplified in the recent work by Borgia et al., where a protein-protein complex with picomolar affinity is shown to lack structure.^2^ While *ΔG*_*bind*_ has been the center of attention of scientific research for decades, little attention has been paid to the factors that determine if a complex will be tight or loose. The source of structural robustness must be sought on sharp (and possibly transitory) energetic barriers that keep the atoms in their positions of equilibrium. Such hypothetical barriers, like the ones that determine binding kinetics, could have their origin in intramolecular (i.e. conformational rearrangement), bimolecular (e.g. repulsive transitional configurations) or many-body effects (e.g. desolvation).^10^ But they will only provide structural stability if the barriers are steep and located very close to the position of minimum energy. In that respect, hydrogen bonds (HBs) are ideal candidates because they have strict distance and angular dependencies^11^ and are one of the most frequent interaction types in protein-ligand complexes.^12^ The contribution of HBs to *ΔG*_*bind*_ has been largely debated in the literature.^13–17^ The current consensus is that it is highly variable and context dependent, but their contribution to thermodynamic stability is 1.8 kcal mol^-1^ at the most.^14^ However, due to desolvation, the transitional penalty of breaking a HB can be much larger.^18^ Indeed, we have shown that this is the case for water-shielded HBs, which can even act as kinetic traps.^19^ More recently, we have also shown that formation of structurally robust intermolecular HBs at specific positions is a necessary condition for binding, and have developed a method to assess the robustness of individual HBs that is very effective in virtual screening applications.^20^

With this background, we decided to perform a systematic investigation of the possible role of HBs as structural anchors of protein-ligand complexes. Our findings not only confirm a general role of HBs as source of structural stability, but also offer a new perspective to understand and design ligand-receptor complexes.

## RESULTS AND DISCUSSION

Using Dynamic Undocking (DUck), an MD-based computational procedure,^20^ we have assessed the robustness of every HB in a set of 79 drug-like protein-ligand complexes from the Iridium Data Set.^21^ Detailed information about the data set and the selection criteria is presented in Supplementary Methods and Supplementary Table 1. Each HB was pulled to a distance of 5 Å, according to the DUck protocol reported previously.^20,22^ In this way, we obtain a work value (*W*_*QB*_) that reflects the cost of breaking each HB. In other words, the W_QB_ value indicates if the interaction under investigation gives rise to a narrow (local) minimum in the free-energy landscape, and estimates its depth. Based on our previous research, we define HBs as robust (i.e. capable of providing structural stability) if *W*_*QB*_ > 6 kcal mol^-1^, labile if *W*_*QB*_ < 4 kcal mol^-1^ and medium otherwise.

The distribution of work values for the entire set of 345 HBs ranges from 0 to 26 kcal mol^-1^, with a of maximum probability in the 0-6 kcal mol^-1^ region and a gradual decrease thereafter (Fig.1a). Noteworthy, more than half HBs (57.4%) are robust. In order to provide a critical assessment of these results, we have sought correlation with experimental observables and have also considered if W_QB_ values might be dominated by the interaction energies. Larger W_QB_ values imply a narrower minimum and, thus, restricted mobility, which should translate into a more localized electron density, that is, lower crystallographic B-factors. As B-factors are heavily influenced by the refinement methods used and their absolute values can be meaningless,^23,24^ we have normalized the B-factor of the ligand atom that makes the hydrogen bond relative to the average B-factor of the whole ligand. Encouragingly, atoms forming HBs with larger W_QB_ values tend to have lower relative B-factors (Supplementary Fig.1). A second aspect to consider is whether DUck calculations merely reflects short-range protein-ligand interaction, or – as intended – it captures a global effect that considers enthalpic and entropic contributions from both the solute and the solvent. Lack of correlation between interaction energies and W_QB_ confirms that the latter is true (Supplementary Fig.2). Of particular interest is to assess the effect of charge reinforcement on HBs, as the energetic, entropic and solvation terms of neutral hydrogen bonds and salt bridges are drastically different.^25^ We have classified all HBs into neutral, mixed (ionic-neutral) and salt bridges (Fig.2, Supplementary Table 2). We find that salt bridges are only very slightly skewed towards more robust interactions than neutral HBs. The distributions were compared with two sample Kolmogorov-Smirnov statistical test, yielding p-value of 0.08. Mixed types are completely indistinguishable from neutral ones (p-value = 0.42). Unexpectedly, the maximal values are equal across all three categories. Theoretically, ionic species could provide even larger energetic barriers because their desolvation costs are much larger. We speculate that there may be no biological use for them, as the maximal *W*_*QB*_ values observed here already ensure very robust and long-lived structures.

The distribution of robust HBs is rather inhomogeneous across complexes, as they have 2.5 on average, but a quarter of the complexes have none (Fig.1b). Considering that structural stability is not a requisite for tight binding and that HBs may not the only mechanism capable of providing structural stability, it is striking that 75% of the complexes in this set are anchored through HBs. A further 14% of complexes present medium values and only in 9 cases (11%) all their HBs are labile (Supplementary Fig.3). Two of those cases are very low affinity complexes. In the remaining cases, structural stability might be provided by other mechanisms or may be lacking (see examples in Supplementary Fig.3). It is important to note that the level of structural stability reported here may be overestimated due to the composition of the data set, entirely derived from X-ray crystallography, a technique that relies on order to solve structures.

**Figure 1.**
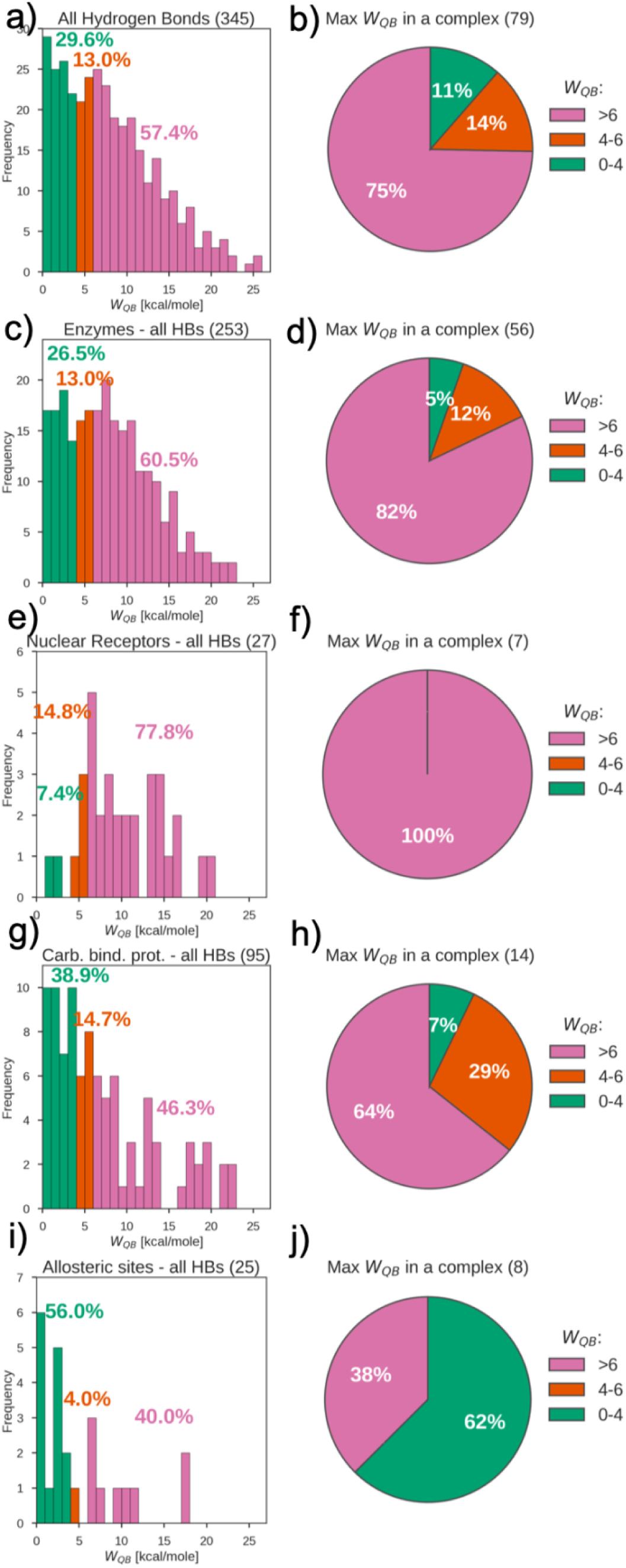
Histograms of frequency of HBs by *W*_*QB*_ value for: a) all simulated HBs (345), c) HBs in enzyme active sites (253), e) HBs in the ligand binding site of nuclear receptors (27), g) HBs in carbohydrate binding sites (95), i) HBs in allosteric sites (25). Pie charts showing share of complexes with at least one robust HB (*W*_*QB*_ > 6 kcal mol^-1^, pink), all labile HBs (*W*_*QB*_ < 4 kcal mol^-1^, green) or intermediate situations (red) for: b) all simulated complexes (79), d) enzymes (56), f) nuclear receptors (7), h) carbohydrate binding site (14), j) allosteric sites (8).

**Figure 2.**
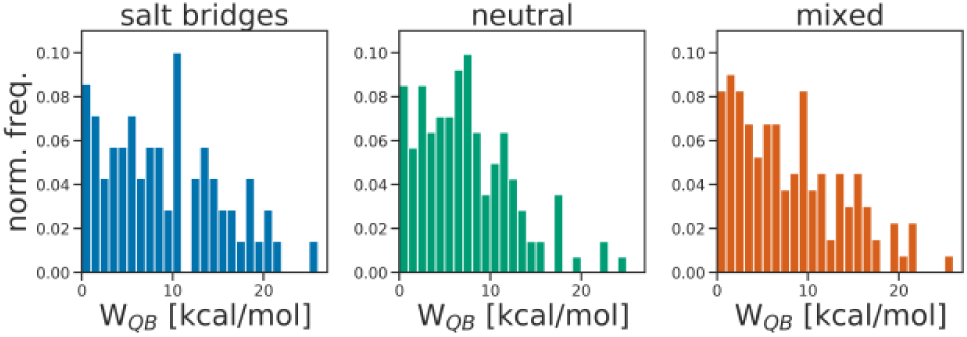
Histograms presenting distribution of *W*_*QB*_ values in the set of HBs from Iridium DS, divided into salt bridges, neutral and mixed (ion-neutral) interactions.

Splitting this analysis by the type of binding site (Fig.1c-j, Supplementary Table 3) provides strong indication that the behavior is dictated by the nature of the receptor. The proportion of robust complexes increases to 82% in the case of enzyme active sites, which speaks about the need of keeping the substrate in place for efficient catalysis. Nuclear receptors form fewer HBs with their ligands, but most of them (78%) are robust and all ligands (100%) are well anchored. In this case, forming a rigid structure may be necessary to stabilize the AF2 co-regulatory protein binding surface in an optimal conformation for co-activator binding.^26^ Carbohydrate binding sites, on the other hand, form many more HBs with their ligands, but a lower proportion of robust ones (46%). Finally, in the case of allosteric ligands, only 40% of complexes are robust, suggesting that these sites tend to yield looser complexes. As demonstrated in the case of HIV reverse transcriptase inhibitors (Fig.3C), lack of robust HBs does not preclude tight binding. In fact, a multiplicity of binding modes might be beneficial to preserve binding affinity when the target is mutated, thus averting resistance.^27,28^ While the distribution of HB strength between the four types of binding sites that we have defined is quite different (see Supplementary Tables 4 and 5 for statistical tests), individual cases can deviate from the norm (e.g. the allosteric ligand 1YV3 is extremely robust) and more examples will be needed to reach firm conclusion about site-dependence.

**Figure 3.**
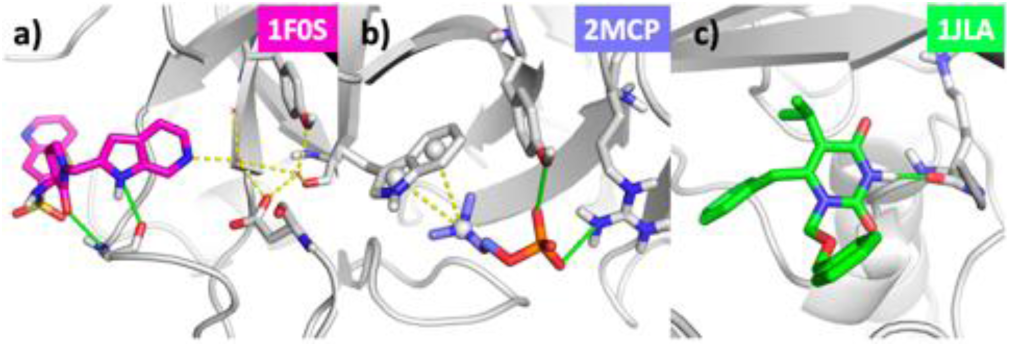
Structures of protein-ligand complexes that form potentially labile structures (all HBs weaker than 4 kcal/mol). Weak hydrogen bonds (W_QB_ < 4 kcal/mol) marked in green. a) Complex of FXa with inhibitor RPR208707 (PDB id 1F0S; *K*_*i*_ = 18 nM) forms two direct, but labile, HBs with the protein. An additional water-mediated HB with the catalytic residues (yellow dotted lines) might provide structural stability. b) An antibody that recognizes phosphocholine (PDB id 2MCP) forms two charge-reinforced but labile HBs. A cation-pi interaction (yellow dotted lines) might provide structural stability. c) Reverse transcriptase inhibitor (PDB id 1JLA; *IC*_*50*_ = 6 nM) forms a single but labile HB with the protein. No other source of structural stability is apparent.

We analyzed the distribution of robust HBs and found that they tend to concentrate on one part of the ligand (Supplementary Fig.4). To better understand this observation, all HBs in each complex were clustered, based on their distance in space, into fragment-sized group of atoms (Supplementary Fig.5). In the majority of complexes (62%) robust HBs were located in a single group, forming a strong structural anchor (Fig.4, Supplementary Table 6). The concentration of robust interactions on a single site, allowing a some degree of movement to the other parts, minimises the entropic costs and can be desirable from a binding affinity perspective.^6^ Only 23% of ligands form two structural anchors on separate regions, though this is more common in the case of carbohydrate-binding proteins (Supplementary Table 7). Three exceptional ligands manage to form 3 distinct stable anchors. Interestingly, they have completely unrelated functions, chemical structures and physical properties but – at least in two of those cases – there is a possible functional explanation for the extreme robustness (Fig.5).

**Figure 4.**
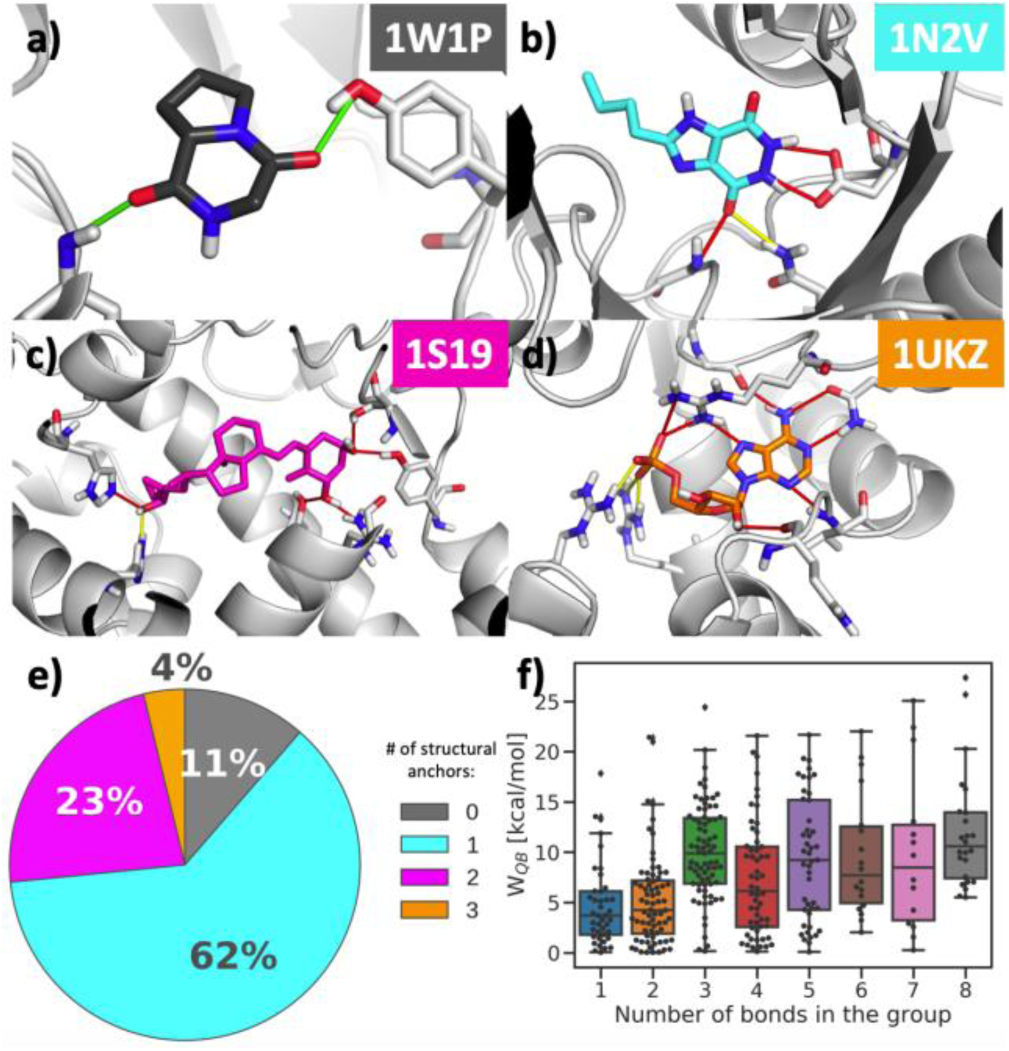
Division of complexes based on the number of structural anchors. Representative of each group is presented in the following image: a) 0 anchors: Chitinase B with inhibitor (PDB id 1W1P; IC_50_ = 5 mM); b) 1 anchor: Queuine tRNA-ribosyltransferase with inhibitor (PDB id 1N2V; *K*_*i*_ = 83 µM), c) 2 anchors: Vitamin D3 receptor with calcipotriol (PDB id 1S19; *K*_*d*_ = 0.31 nM) and d) 3 anchors: Uridylate kinase - AMP (PDB id 1UKZ). e) Pie chart presenting distribution of number of anchors across the data set. f) Distribution of strength of HBs (W_QB_) versus the number of HBs per group of atoms. Weak hydrogen bonds (W_QB_ < 4 kcal/mol) marked in green, medium (4 ≤ W_QB_ < 6 kcal/mol) in yellow and strong (W_QB_ ≥ 6 kcal/mol) in red.

**Figure 5.**
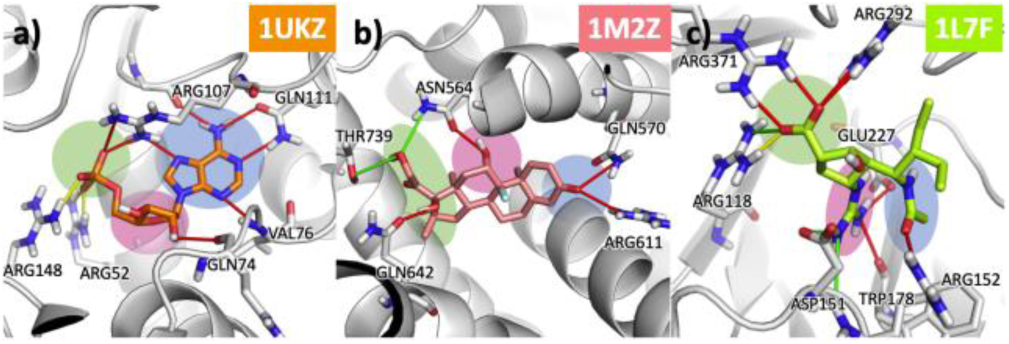
Structures of complexes with three binding anchors (shaded areas). a) Uridilate kinase with AMP (PDB id 1UKZ) where the base, ribose and phosphate of the nucleotide are forming three distinctive centres of interactions. Structural stability may be necessary for efficient catalysis. b) Glucocorticoid receptor ligand-binding domain bound to dexamethasone (PDB id 1M2Z; *K*_*d*_ = 19 nM). The ligand has three regions that form robust interaction, well separated in space but located on the steroid core, thus behaving as a single rigid block. Structural stability may be necessary for agonistic response. c) Influenza virus neuraminidase with inhibitor BCX-1812 (PDB id 1L7F; *K*_*i*_ single digit nM for various virus strains). Three different functional groups branching out of the pentane scaffold form robust interactions in this extremely polar and solvent exposed binding site. Weak hydrogen bonds (W_QB_ < 4 kcal/mol) marked in green, medium (4 ≤ W_QB_ < 6 kcal/mol) in yellow and strong (W_QB_ ≥ 6 kcal/mol) in red.

The distribution of *W*_*QB*_ per number of HBs in a local group (Fig.4f) is suggestive of cooperative behavior. HBs in isolation usually do not form robust interactions (mean and median values: (4.7 ± 4.1) and 3.7 kcal mol^-1^, respectively), although in exceptional cases they can reach values above 10 kcal mol^-1^. By contrast, when three or more HBs cluster together, formation of robust complexes is the most common outcome (mean and median values: (9.4 ± 5.8) and 9.0 kcal mol^-1^, respectively). The HBs within these clusters present relatively similar W_QB_ values (Supplementary Fig.6), suggesting that they often behave in a concerted-like manner. This synergic and mutually dependent behavior not only ensures higher barriers to dissociation, but is also well-suited to provide selectivity, as small changes in the composition or geometry of one of the partners may result in large changes in magnitude of W_QB_ (see example in Supplementary Fig.7).

The observation that most drug-like ligands combine tightly-bound regions with looser makes us wonder about fragment-sized ligands. Do they valance order and disorder in some other way (e.g. using fewer attachment points)? Or, perhaps, depending on the site they bind to, they are either dynamic or fully constrained? In order to answer these questions, we have extended our analysis with a set of 27 fragment-protein complexes (126 individual HBs) from the SERAPhiC dataset.^29^ Strikingly, we find that fragments have an almost identical behavior to standard ligands, with 49% of robust HBs (2.3 per ligand) and 73% of ligands presenting at least one robust interaction. The distribution and maximal W_QB_ values are also very similar (Fig.6). This indicates that, proportionally, fragments are more static than standard ligands. This agrees with the observations that fragments have a more enthalpic binding^30^ and that they have a higher proportion of buried HBs.^31^ It also justifies that, in spite of their low binding affinity, most fragments already have a well-defined binding mode that serves as a foundation from which to spread and catch additional interactions. However, not all fragments form robust interactions and we propose that these are less suitable as starting points because their binding mode can change, confounding structure-activity interpretation and rendering optimization more difficult. Indeed, fragments are known to change their binding mode when evolved into larger molecules.^7,32–36^ These may be attempts at building on what is assumed to be a solid foundation but turns out to be unstable ground, a possibility that we shall investigate in the future. It should also be noted that the fraction of well-anchored fragments may be different for fragments hits that fail to crystallize. The overlap between X-ray crystallography and other biophysical screening methods can be rather low^37^ and progressing fragments that fail to crystallize is deemed difficult but worthwhile.^38^

**Figure 6.**
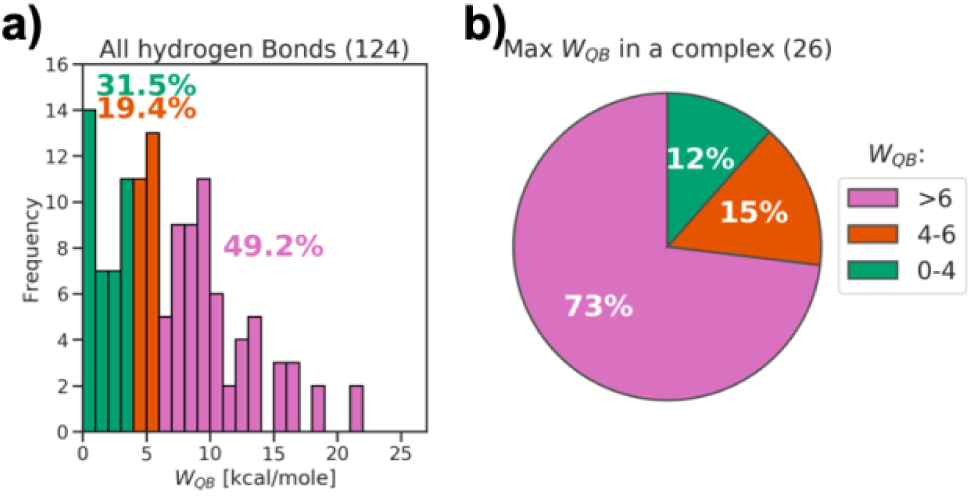
a) Histograms of frequency of HBs by *W*_*QB*_ value for all simulated HBs (124) in SERAPhiC dataset. Pie charts showing share of complexes with at least one robust HB (*W*_*QB*_ > 6 kcal mol^-1^, pink), all labile HBs (*W*_*QB*_ < 4 kcal mol^-1^, green) or intermediate situations (red) for all protein-fragment complexes (26).

Finally, we want to consider what is the origin of the free energy barrier that causes structural stability. Knowing that a HB has a large *W*_*QB*_ value can be likened to knowing the *k*_*off*_ of a compound without knowing the *k*_*on*_ nor *ΔG*_*bind*_: larger values may indicate that it has a higher transition state (if *ΔG*_*bind*_ remains the same; Fig.7a), that the complex is thermodynamically more stable (if *k*_*on*_ remains the same; Fig.7b), or a combination thereof. In this data set, we find that anchoring sites often correspond to binding hot spots. This is indeed the case for all kinases and proteases, which have a well-known binding hot spot (Supplementary Table 3, Supplementary Fig.5), as well as for most fragments. In such cases, *ΔG*_*bind*_ must be a component of *W*_*QB*_, but there is no correlation between both magnitudes (Supplementary Fig.8), as already noted.^20^ Thus, we conclude that *W*_*QB*_ must be largely dominated by a transitory dissociation penalty. The origin of this penalty can be explained by a physical decoupling between HB rupture and resolvation, as described for water-shielded hydrogen bonds.^19^ In support of this view, several studies of the reverse event have identified desolvation of the binding pocket as the rate-limiting step in ligand association.^18,39,40^ Indeed, solvent exposed HBs invariably lead to low *W*_*QB*_ values (but note that they can be thermodynamically stable),^41^ whereas water-shielding is a necessary but not sufficient condition of robust HBs (Supplementary Fig.9).

**Figure 7.**
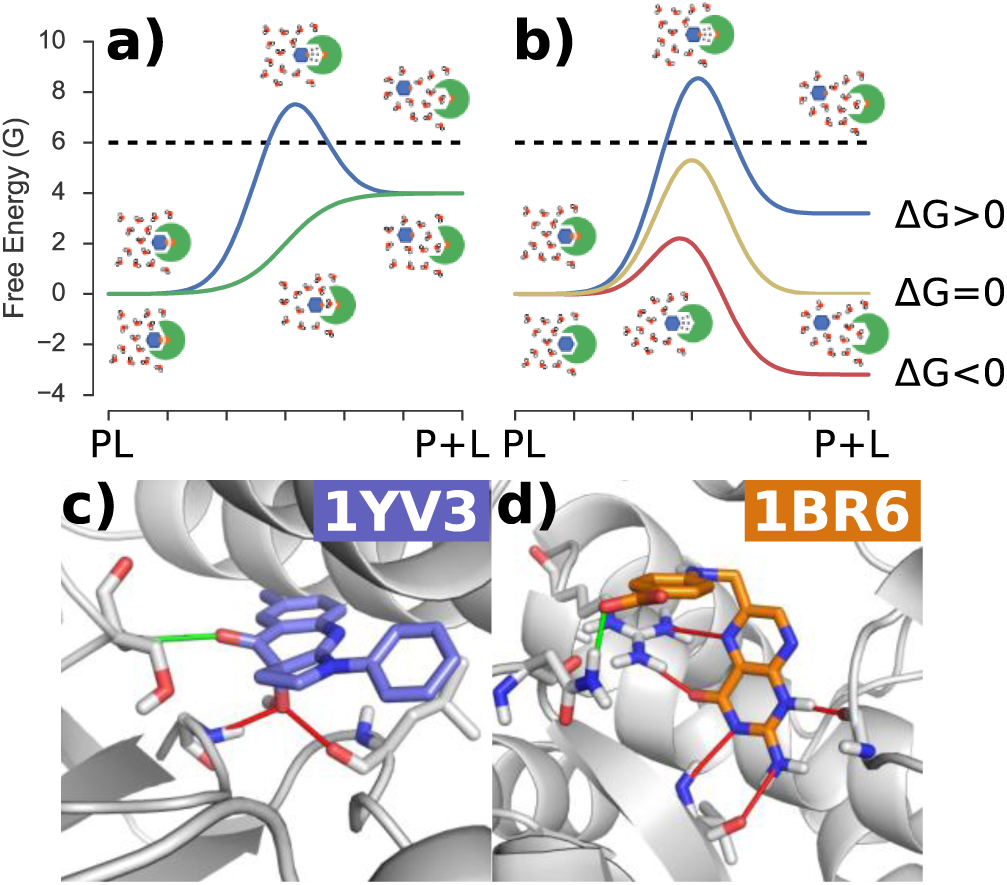
Ways of achieving structural robustness. a) Idealized representation of two dissociation pathways for complexes with the same *ΔG*_*bind*_ and different desolvation costs. The images above the blue curve shows the state of the system in bound, transition and unbound state of complex with well shielded, stable hydrogen bond. The images below the green curve show analogous images for the complex with solvent exposed hydrogen bond. b) Likewise for two complexes with the same desolvation cost but different *ΔG*_*bind*_. The images represent complexes with excellent shape complementarity that form (above the blue curve) or don’t form (below the red curve) favorable hydrogen-bonding pairs. The black dashed line marks the energy cutoff that classifies bond as structurally stable. c) Example of a complex with high dissociation cost due to extreme water-shielding. d) Example of a complex with high dissociation cost due to a tight network of multiple HBs. Weak hydrogen bonds (W_QB_ < 4 kcal/mol) marked in green and strong (W_QB_ ≥ 6 kcal/mol) in red.

## CONCLUSION

Taken together, our results show that structural stability is a common property of protein-ligand complexes, but not an universal one. Cases of loose complexes, while relatively rare (10-20%), can be found even in a dataset originating exclusively from X-ray crystallography, a technique that requieres structural homogeneity of the sample. The proportion could be larger amongst ligands that fail to crystallize. The level of residual mobility is also larger and more common than the static X-ray structures lead to think, as also concluded by a recent independent study.^4^ In fact, most complexes balance order and disorder by combining a firm anchor with more relaxed peripheral interactions. Depending on the nature of the ligand and the binding site, each complex adopts a particular degree of robustness, that ranges from the very tight (e.g. nuclear receptor agonists) to the very loose (e.g. HIV-RT allosteric inhibitors). Each one of these solutions entails important consequences that have, so far, been neglected in drug design. First of all, a firm anchor provides a framework from which to grow and capture additional interactions, and the preservation of a common binding mode helps interpreting structure-activity relationships. This is particularly important for fragments as starting points for lead discovery. Secondly, structural robustness can have functional implications, particularly in the case of receptors, where flexibility has been linked to the agonist/antagonist response.^26,42^ Thirdly, structural stability implies an entropic penalty and must be balanced to avoid loss of potency.^6,43^ Finally, the deep and narrow energetic minima that cause rigidity also imply large penalties for small recognition defects, thus increasing the fidelity of the recognition event. This has been shown for protease-substrate pairs^44^ and HIV-protease inhibitors.^45^ In conclusion, this work opens up the possibility of understanding and designing structural robustness in ligand-receptor complexes. We suggest that robustness analysis, which can help understand and control the level of mobility, should be an essential part of ligand design, not least because rigid parts demand more precise complementarity than flexible ones. Qualitatively, a visual inspection can reveal water-shielded HBs (Fig.7c) and HB clusters (Fig.7d), which are tell-tale signs of robustness. Quantitatively, DUck simulations offer an inexpensive and automated protocol to calculate *W*_*QB*_. While HBs appear to be the most common means of achieving structural robustness, other interaction types (e.g. cation-pi, water-mediated HBs, halogen bonds) should be considered in the future.

## Supporting information

Iridium_structures

SERAPhiC_structures

## ASSOCIATED CONTENT

### Supporting Information

Detailed methods, additional figures and tables. Set of sdf files containing ligands and mol2 files containing protein structures.

## ACKNOWLEDGMENT

We thank C. Galdeano, F. J. Luque and the anonymous reviewers for helpful discussions and manuscript revision. The research was funded under the EU Horizon 2020 programme, Marie Sklodowska-Curie grant agreement No. 675899 (FRAGNET). Spanish Ministerio de Economia (SAF2015-68749-R). Catalan government (2014 SGR 1189). Barcelona Supercomputing Center (computational resources).

## AUTHOR CONTRIBUTIONS

X.B. designed the project; M.M. performed calculations; S.R.C. contributed new analytic tools; M.M. and X.B. analysed data and wrote the manuscript.

