## Supplementary Information for "Are protein-ligand complexes robust structures?"

### Supplementary Methods

#### Data Sets

##### Iridium

As an object of calculations we selected a set of 120 high-trustworthy structures of protein-ligand complexes collected in Iridium Data set.[1] Refined protein structures and binding modes of ligands with assigned protonation and tautomeric states were provided by developers of the set. Due to limitations of used methods, some cases were excluded from calculations. First excluded group were complexes that did not possess any intermolecular hydrogen bonds (3 cases in Iridium). Coordination complexes with metal ions (28 cases) were also excluded. They are known to be stable structures, however they are not faithfully represented by the molecular mechanics force-field. Cases of multiple ligands in a binding pocket (5 cases) were also excluded, due to complexity of analysis, as well as ligands that were deemed non-drug like (6 cases). In case of 1hgh ligand was present in two binding pockets, differing in affinity.[2] In this case we simulated and analyzed both of the pockets separately. After the reduction of the set we were left with 78 complexes and 79 binding pockets. Classification of the structures is presented in Supplementary Table 1.

##### SERAPhiC

Second validation consisted of 53 high-quality X-ray models of fragment-protein complexes.[3] Protonation states of both protein structures and ligands were provided by developers of the data set. The same as in case of Iridium, some systems were excluded from the simulations: no hydrogen bonds (7 cases), additional ligands (5 cases) and metal ions (17 cases). After the reduction, 26 binding pockets were left in 24 different PDB structures. Three structures (1e2i, 1ofz, 2hdq) possess two ligands per protein indicated by "a" and "b" next to PDBid. Both pockets in entries 1ofz and 2hdq were treated as separate cavities in simulation and analysis stage. Complex 1e2i possesses 2 enantiomers in the same binding pocket. Both isomers were simulated separately, but in the analysis both of them were treated as a single pocket. Classification of the structures is presented in Supplementary Table 8.

#### Systems preparation

Structures of protein ligand complexes were provided by Iridium Data set's developers. The ligands had preassigned protonation and tautomeric states, which were then verified to reassure the right ligand parametrization. The protein structures were initially prepared by protonation at pH = 7 with MOE.[4] The protonation states were then inspected visually and questionable cases were verified with publication associated with PDB code.

Hydrogen bonds in complexes were identified using Chimera FindHBond function.[5] Then each bond was treated as a core for the individual simulation system. In order to perform Dynamic Undocking only a part of the protein was selected for the MD simulation. So-called chunk consists of residues necessary to preserve ligand's key interaction. First, crystal water molecules, additional molecules and ions were removed from protein structure. Then all residues with at least one atom within 7 Å of the key interaction were selected. Additional residues were included in order to prevent formation of artificial solvent channels. The residues that were not selected, were removed from the structure, forming the base chunk. In order to remove artificial charges, truncated chains were then acetylated or N-methylated, as needed. Reducing the system minimizes the influence of peripheral interactions and simplifies dissociation pathway. Additionally, it reduces time of calculation.

### Dynamic Undocking

469 (345 for Iridium and 124 for SERAPhiC) individual hydrogen bonds were identified for 105 binding pockets for each bond individual system was created containing ligand and chunk of a protein. Then each system was prepared according to Dynamic Undocking protocol,[6,7] using parameters:

- Equilibration length: 1 ns
- MD chunk length: 0.5 ns
- SMD length: 0.5 ns
- SMD displacement: 2.5 Å
- Force constant: 50 kcal $mol^{-1}$ $Å^{-2}$
- $W_{QB}$ threshold: 0.4 kcal $mol^{-1}$ (to avoid early termination of the protocol)
- Maximum DUck SMD runs: 19

Preparation of the simulations was run by previously developed Molecular Operating Environment[4] Scientific Vector Language script available online. The script performs the following actions:

- Calculates AM1-BCC charges for the ligand.[8]
- Assigns atom types and non-bonded parameters to the ligand, using parm@Frosst force field.[9]
- Identifies the pair of atoms making the key interaction.
- Writes the files necessary to carry out the MD simulations with AMBER 12.[10] (AMBER99 force field is used to parametrize protein).
- Generates valid topology and coordinate files for each system using AMBER's tLeap.

With those parameters each bond was subjected to 40 steered molecular dynamics (SMD) trajectories at 300 K and 325 K (20 trajectories in each temperature). In case of low energy bonds, the simulations were terminated after reaching the value below 0.4 kcal $mol^{-1}$. Setting the early termination threshold allows avoiding simulation of low stability systems. In this project, the value is very low and does not influence many systems. Here, only 4 systems in Iridium and 2 in SERAPhiC were terminated before reaching full 40 SMDs. All the simulations were performed with AMBER14 adapted for running in graphics processing unit (GPUs) and executed at the Barcelona Supercomputing Centre using NVIDIA Tesla M2090 GPUs. The work necessary to reach quasi-bound state ($W_{QB}$) in which the hydrogen bond has just been broken, was calculated for each individual trajectory.[6] The lowest value out of the set was then selected as a final stability score, because in that trajectory the ligand followed dissociation pathway that allowed it to escape the pocket. The final $W_{QB}$ value reflects bond's resistance to deviate from the optimal geometry. All the bonds are classified with their location, stability ($W_{QB}$), character (salt bridge, charged or neutral) in the Supplementary Tables 2 and 9.

It is worth noticing that individual $W_{QB}$ values do not have an associated error value. To assess the convergence of simulation, 40 SMDs per hydrogen bond were randomly split into 4 equal parts of 10 SMDs. For each part the lowest $W_{QB}$ value was identified and then 4 selected values were averaged into a new value, called "Mean-$W_{QB}$" with calculated standard deviation (STD). Both of values are included in Supplementary Tables 2 and 9. The conventionally calculated $W_{QB}$ and Mean-$W_{QB}$ are very similar (Supplementary Fig. 11), with correlation equal to 0.99.

***Supplementary Table 1*** *Classification of complexes from Iridium DS*

| | |
|---|---|
| Simulated structures | 1a28, 1ai5, 1b9v, 1br6, 1c1b, 1cvu, 1dds, 1exa, 1ezq, 1f0s, 1f0t, 1f0u, 1fcx, 1fcz, 1fh8, 1fh9, 1fhd, 1fm6, 1fvt, 1g9v, 1gm6, 1h1p, 1h1s, 1hgh-1, 1hgh-2, 1hgi, 1hgj, 1hwi, 1ivb, 1ivd, 1ive, 1ivf, 1jla, 1k1j, 1k3u, 1ke5, 1l2s, 1l7f, 1lpz, 1lqd, 1m2z, 1ml1, 1mq6, 1mts, 1n2j, 1n2v, 1n46, 1of1, 1owe, 1oyt, 1pmn, 1q1g, 1q41, 1qhi, 1rob, 1s19, 1tow, 1tt1, 1u4d, 1ukz, 1ulb, 1unl, 1uou, 1v0p, 1w1p, 1w2g, 1x8x, 1ydr, 1yds, 1ydt, 1yv3, 1yvf, 1ywr, 2ack, 2br1, 2mcp, 2pcp, 3ptb, 4ts1 |
| No HB | 1ctr, 1fl3, 1p2y |
| Not drug like | 1fjs, 1fm9, 1fq5, 1gwx, 1hgg, 1pso |
| Metal ion | 1azm, 1cx2, 1dd7, 1eoc, 1frp, 1hdy, 1hq2, 1hww, 1iy7, 1jd0, 1lrh, 1mbi, 1mmv, 1mzc, 1n1m, 1oq5, 1p62, 1r58, 1r9o, 1uml, 1xm6, 1xoq, 1yqy, 2ctc, 2tmn, 4aah, 4cox, 1hp0 |
| Additional ligand | 1d3h, 1hnn, 1hvy, 1pbd, 1sq5 |

***Supplementary Figure 1*** *Normalized β-factor of atom that makes the hydrogen bond in function of calculated $W_{QB}$. β-factor has been normalized against β-factors of other atoms of the ligand in PDB structure. Liable hydrogen bonds are depicted in blue and strong hydrogen bonds are depicted in green. The atoms with β-factor below the average are marked in magenta and the ones with β-factor above the average are marked in red. The side panels show the distribution of selected points as a kde-plots.*

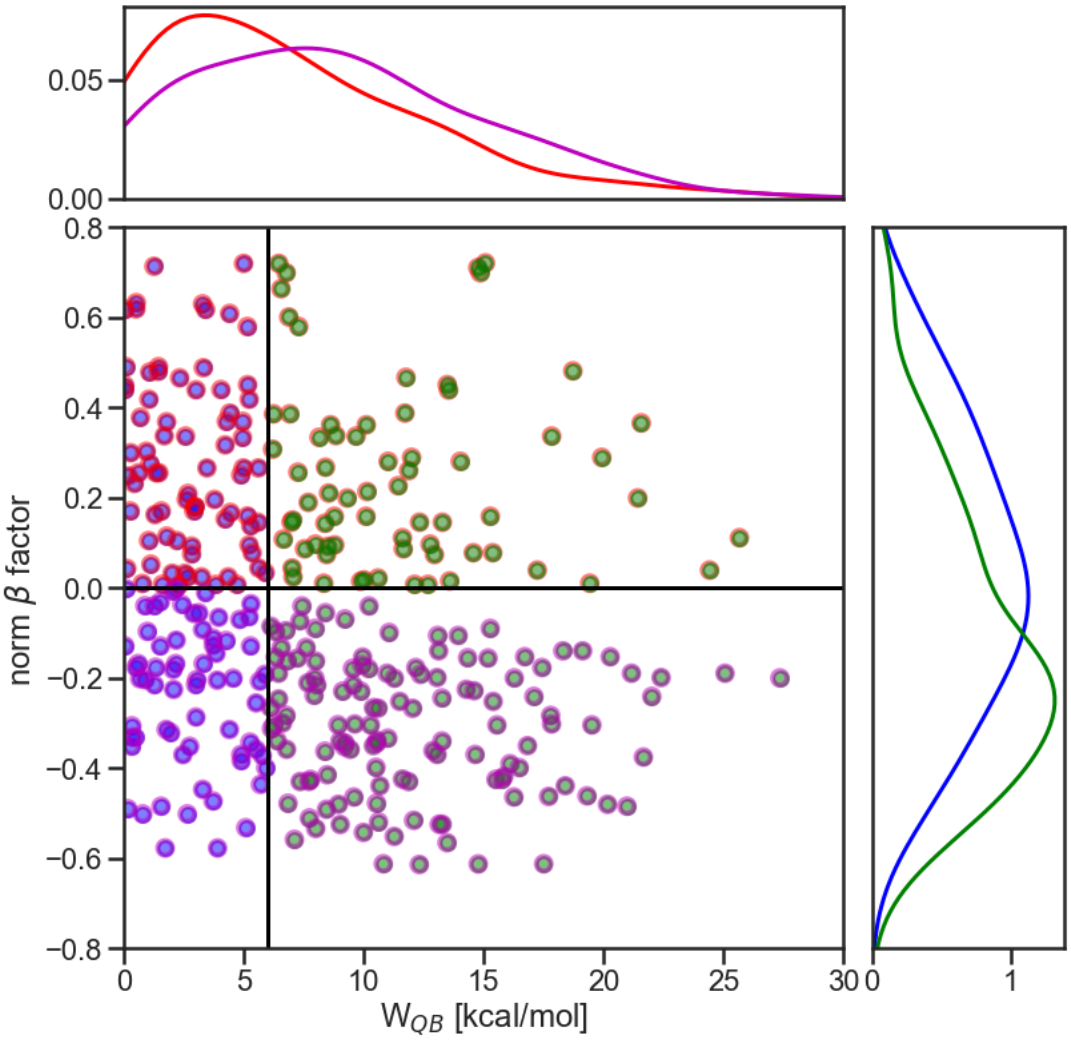

***Supplementary Figure 2*** *A series of plots comparing $W_{QB}$ value with potential energy calculations for hydrogen bonds from Iridium DS. The energy was computed as electrostatic potential energy (plots in the middle), and sum of electrostatic and Van der Waals potential energy (right plots). The energies were plotted for: a) interaction between atoms that form hydrogen bond, b) interaction of atom from ligand that form hydrogen bond with all atoms from chunk, c) interaction of atom form chunk that from hydrogen bond with all atoms from ligand, and d) Interaction of atoms that form hydrogen bond with the rest of the structure.*

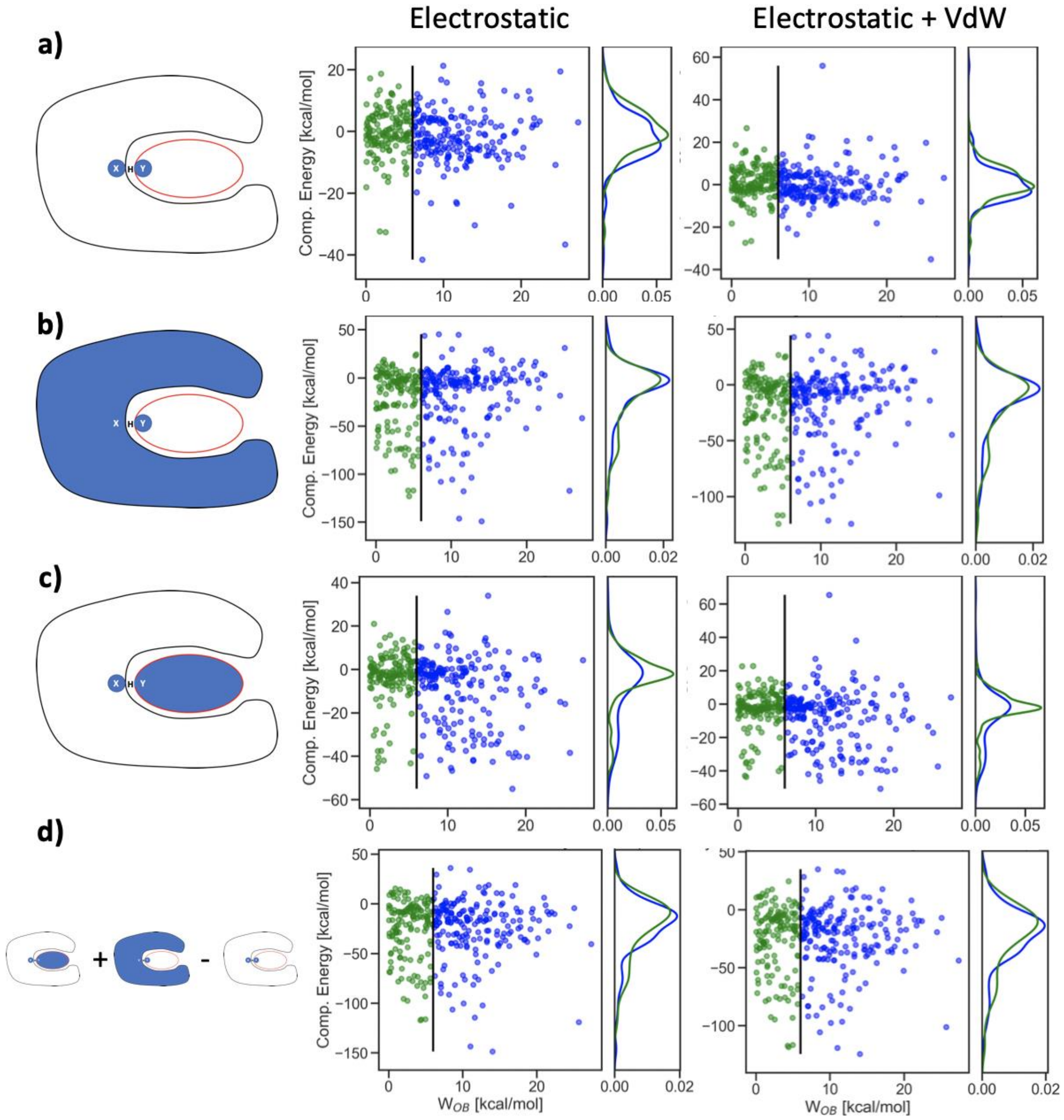

***Supplementary Table 2** List of hydrogen bonds for simulated systems from Iridium DS. The bonds are classified by the complex's PDB code, residue number and atom name of protein's atom that makes the hydrogen bond. Table additionally contains, stability evaluation in a form of calculated $W_{QB}$ value, Mean_$W_{QB}$ value with standard deviation, the orientation of hydrogen bond (donor/acceptor function of protein's atom), and the bond character (information if the bond makes a salt bridge or neutral interaction).*

| PDB | RESIDUE | PROT_ATOM | # SMD | $W_{QB}$ [KCAL/MOL] | MEAN_WQB [KCAL/MOL] | STD [KCAL/MOL] | PROT (D/A) | SALT BRIDGE | NEUTRAL HB |
|---|---|---|---|---|---|---|---|---|---|
| **1A28** | ARG766 | NH2 | 40 | 4.99 | 5.37 | 0.22 | D | 0 | 0 |
| | GLN725 | NE2 | 40 | 6.46 | 7.19 | 0.59 | D | 0 | 1 |
| **1AI5** | ASN241 | ND2 | 40 | 5.79 | 6.03 | 0.27 | D | 0 | 0 |
| | SER1 | OG | 40 | 7.43 | 8.99 | 1.13 | D | 0 | 0 |
| **1B9V** | ARG116 | NH2 | 6 | 0.32 | 0.65 | 0.24 | D | 1 | 0 |
| | ARG150 | NH1 | 40 | 6.15 | 6.42 | 0.19 | D | 0 | 0 |
| | ARG292 | NH2 | 40 | 0.16 | 2.58 | 1.56 | D | 1 | 0 |
| | ARG374 | NH2 | 40 | 2.75 | 3.82 | 0.71 | D | 1 | 0 |
| | GLU276 | OE2 | 40 | 0.66 | 1.10 | 0.29 | A | 0 | 0 |
| | TRP177 | O | 40 | 4.87 | 5.84 | 0.75 | A | 0 | 1 |
| **1BR6** | ARG180 | NH1 | 40 | 21.67 | 21.97 | 0.19 | D | 0 | 0 |
| | ARG180 | NH2 | 40 | 16.10 | 16.27 | 0.14 | D | 0 | 0 |
| | ASN78 | ND2 | 40 | 1.43 | 1.79 | 0.23 | D | 0 | 0 |
| | GLY121 | O | 40 | 11.62 | 12.88 | 0.93 | A | 0 | 1 |
| | VAL81 | N | 40 | 17.71 | 19.66 | 1.19 | D | 0 | 1 |
| | VAL81 | O | 40 | 19.31 | 20.23 | 0.87 | A | 0 | 1 |
| **1C1B** | LYS101 | O | 40 | 2.82 | 3.33 | 0.49 | A | 0 | 1 |
| **1CVU** | SER530 | OG | 40 | 1.95 | 2.21 | 0.27 | D | 0 | 0 |
| | TYR385 | OH | 40 | 5.08 | 7.67 | 2.79 | D | 0 | 0 |
| **1DDS** | ASH27 | OD1 | 40 | 11.01 | 12.21 | 0.76 | A | 0 | 1 |
| | ASH27 | OD2 | 40 | 10.55 | 10.80 | 0.31 | D | 0 | 1 |
| | ILE5 | O | 40 | 4.40 | 5.39 | 0.61 | A | 0 | 1 |
| | ILE94 | O | 40 | 1.76 | 2.29 | 0.39 | A | 0 | 1 |
| **1EXA** | ARG278 | NH1 | 40 | 6.76 | 6.95 | 0.18 | D | 1 | 0 |
| | LEU271 | O | 40 | 13.26 | 13.36 | 0.07 | A | 0 | 1 |
| | MET272 | SD | 40 | 6.32 | 7.71 | 0.81 | A | 0 | 1 |
| | SER289 | N | 40 | 8.41 | 9.17 | 0.50 | D | 0 | 0 |
| | SER289 | OG | 40 | 14.86 | 15.65 | 0.53 | D | 0 | 0 |
| **1EZQ** | ASP189 | OD1 | 40 | 9.98 | 11.07 | 1.03 | A | 1 | 0 |
| | ASP189 | OD2 | 40 | 10.40 | 13.14 | 2.06 | A | 1 | 0 |
| | GLU97 | O | 40 | 1.78 | 2.60 | 0.69 | A | 0 | 0 |
| | GLY219 | N | 40 | 3.95 | 4.11 | 0.12 | D | 0 | 1 |
| | GLY219 | O | 40 | 16.83 | 18.10 | 0.86 | A | 0 | 0 |
| **1F0S** | GLY219 | N | 40 | 2.95 | 3.70 | 0.47 | D | 0 | 1 |

| | | | | | | | | | |
|---|---|---|---|---|---|---|---|---|---|
| | GLY219 | O | 40 | 2.91 | 4.00 | 0.77 | A | 0 | 1 |
| **1F0T** | ASP189 | OD1 | 40 | 12.08 | 13.64 | 0.93 | A | 1 | 0 |
| | ASP189 | OD2 | 40 | 18.41 | 20.56 | 1.27 | A | 1 | 0 |
| | GLY219 | N | 40 | 2.42 | 2.76 | 0.32 | D | 0 | 1 |
| | GLY219 | O | 40 | 10.68 | 12.51 | 1.30 | A | 0 | 0 |
| **1F0U** | ASN97 | O | 40 | 0.50 | 1.35 | 0.65 | A | 0 | 0 |
| | ASP189 | OD1 | 40 | 14.64 | 15.34 | 0.42 | A | 1 | 0 |
| | ASP189 | OD2 | 40 | 15.56 | 16.49 | 0.55 | A | 1 | 0 |
| | GLN192 | N | 40 | 1.60 | 2.15 | 0.35 | D | 0 | 1 |
| | GLY219 | N | 40 | 2.30 | 3.12 | 0.62 | D | 0 | 1 |
| | GLY219 | O | 40 | 19.52 | 19.98 | 0.45 | A | 0 | 0 |
| | SER190 | OG | 40 | 13.06 | 14.07 | 0.81 | A | 0 | 0 |
| **1FCX** | MET272 | SD | 40 | 5.29 | 5.71 | 0.25 | A | 0 | 1 |
| | SER289 | N | 40 | 6.65 | 7.63 | 0.93 | D | 0 | 0 |
| | SER289 | OG | 40 | 15.08 | 15.31 | 0.23 | D | 0 | 0 |
| **1FCZ** | SER289 | N | 40 | 8.50 | 9.39 | 0.55 | D | 0 | 0 |
| | SER289 | OG | 40 | 14.76 | 14.85 | 0.06 | D | 0 | 0 |
| **1FH8** | ASN44 | ND2 | 40 | 8.84 | 9.28 | 0.30 | D | 0 | 1 |
| | GLN203 | OE1 | 40 | 13.13 | 13.37 | 0.15 | A | 0 | 0 |
| | GLU127 | OE2 | 40 | 19.13 | 19.74 | 0.40 | A | 1 | 0 |
| | GLU233 | OE1 | 40 | 18.31 | 20.54 | 1.50 | A | 1 | 0 |
| | GLU43 | OE2 | 40 | 8.45 | 8.86 | 0.34 | A | 0 | 0 |
| | HID80 | NE2 | 40 | 17.43 | 18.15 | 0.45 | A | 0 | 1 |
| | LYS47 | NZ | 40 | 12.17 | 14.06 | 1.15 | D | 0 | 0 |
| | TRP273 | NE1 | 40 | 12.95 | 13.80 | 0.50 | D | 0 | 1 |
| **1FH9** | ASN126 | ND2 | 40 | 8.33 | 9.26 | 0.64 | D | 0 | 1 |
| | ASN44 | ND2 | 40 | 8.54 | 8.93 | 0.35 | D | 0 | 1 |
| | GLN203 | OE1 | 40 | 6.55 | 6.69 | 0.11 | A | 0 | 1 |
| | GLU127 | OE2 | 40 | 18.72 | 19.78 | 0.70 | A | 1 | 0 |
| | GLU233 | OE2 | 40 | 19.43 | 20.51 | 1.09 | A | 0 | 0 |
| | GLU43 | OE2 | 40 | 7.99 | 8.60 | 0.48 | A | 0 | 0 |
| | HID80 | NE2 | 40 | 22.01 | 22.51 | 0.29 | A | 0 | 1 |
| | LYS47 | NZ | 40 | 17.10 | 17.77 | 0.42 | D | 0 | 0 |
| | TRP273 | NE1 | 40 | 12.77 | 13.79 | 0.65 | D | 0 | 1 |
| **1FHD** | ASN126 | ND2 | 40 | 10.97 | 11.15 | 0.19 | D | 0 | 1 |
| | ASN44 | ND2 | 40 | 7.26 | 7.75 | 0.43 | D | 0 | 1 |
| | GLN203 | NE2 | 40 | 6.45 | 7.81 | 0.79 | D | 0 | 1 |
| | GLU127 | OE2 | 40 | 7.24 | 7.75 | 0.45 | A | 1 | 0 |
| | GLU233 | OE1 | 40 | 25.07 | 25.55 | 0.73 | A | 0 | 0 |
| | GLU233 | OE2 | 40 | 21.17 | 22.45 | 1.39 | A | 0 | 0 |
| | GLU43 | OE2 | 40 | 5.16 | 6.45 | 1.33 | A | 0 | 0 |
| | HID80 | NE2 | 40 | 22.40 | 22.93 | 0.33 | A | 0 | 1 |

| | | | | | | | | | |
|---|---|---|---|---|---|---|---|---|---|
| | LYS47 | NZ | 40 | 13.05 | 13.96 | 0.62 | D | 0 | 0 |
| | TRP273 | NE1 | 40 | 13.48 | 13.72 | 0.16 | D | 0 | 1 |
| **1FM6** | HIE323 | NE2 | 40 | 1.27 | 2.73 | 1.34 | D | 0 | 1 |
| | HIE449 | NE2 | 40 | 5.22 | 6.91 | 1.01 | D | 0 | 1 |
| | SER289 | OG | 40 | 2.99 | 3.50 | 0.38 | D | 0 | 1 |
| | TYR473 | OH | 40 | 10.60 | 10.97 | 0.28 | A | 0 | 1 |
| **1FVT** | ASP86 | N | 40 | 2.97 | 4.01 | 0.63 | D | 0 | 1 |
| | GLU81 | O | 40 | 4.89 | 5.53 | 0.49 | A | 0 | 1 |
| | LEU83 | N | 40 | 11.87 | 11.96 | 0.12 | D | 0 | 1 |
| **1G9V** | LYS99 | NZ | 40 | 0.93 | 1.19 | 0.19 | D | 0 | 0 |
| **1GM8** | SER1 | OG | 40 | 4.13 | 5.22 | 0.66 | D | 0 | 1 |
| **1H1P** | GLU81 | O | 40 | 11.27 | 11.50 | 0.15 | A | 0 | 1 |
| | LEU83 | N | 40 | 7.74 | 8.45 | 0.51 | D | 0 | 1 |
| | LEU83 | O | 40 | 8.45 | 9.37 | 0.67 | A | 0 | 1 |
| **1H1S** | ASP86 | N | 40 | 1.68 | 2.86 | 0.76 | D | 0 | 1 |
| | ASP86 | OD2 | 40 | 3.01 | 3.38 | 0.22 | A | 0 | 0 |
| | GLU81 | O | 40 | 10.51 | 11.27 | 0.56 | A | 0 | 1 |
| | LEU83 | N | 40 | 11.06 | 11.28 | 0.17 | D | 0 | 1 |
| | LEU83 | O | 40 | 13.59 | 13.71 | 0.13 | A | 0 | 1 |
| **1HGH-1** | ASN137 | N | 40 | 1.08 | 2.29 | 0.81 | D | 0 | 0 |
| | GLU190 | OE2 | 40 | 1.46 | 4.50 | 1.83 | A | 0 | 0 |
| | GLY135 | O | 40 | 3.73 | 4.39 | 0.48 | A | 0 | 1 |
| | HIE183 | NE2 | 40 | 1.34 | 2.31 | 0.57 | D | 0 | 1 |
| | SER136 | OG | 40 | 4.22 | 4.88 | 0.61 | D | 0 | 0 |
| | SER228 | OG | 40 | 0.61 | 1.47 | 0.66 | D | 0 | 1 |
| | TYR98 | OH | 40 | 4.44 | 4.60 | 0.14 | D | 0 | 1 |
| **1HGH-2** | ARG269 | NH2 | 40 | 0.28 | 0.83 | 0.34 | D | 0 | 0 |
| | GLH89 | OE2 | 40 | 0.08 | 0.50 | 0.29 | D | 0 | 1 |
| | GLU72 | N | 40 | 0.04 | 0.84 | 0.48 | D | 0 | 1 |
| **1HGI** | ASN137 | N | 40 | 3.84 | 4.12 | 0.16 | D | 0 | 0 |
| | GLU190 | OE1 | 40 | 3.30 | 3.81 | 0.34 | A | 0 | 0 |
| | GLY135 | O | 40 | 5.15 | 6.17 | 1.04 | A | 0 | 1 |
| | HIE183 | NE2 | 40 | 0.59 | 3.11 | 1.83 | D | 0 | 1 |
| | SER136 | OG | 40 | 3.28 | 5.06 | 1.55 | D | 0 | 0 |
| | SER145 | OG | 40 | 0.69 | 0.91 | 0.18 | D | 0 | 1 |
| | SER228 | OG | 40 | 2.16 | 2.77 | 0.40 | D | 0 | 1 |
| | TYR98 | OH | 40 | 5.77 | 6.69 | 0.96 | D | 0 | 1 |
| **1HGJ** | ASN137 | N | 40 | 1.12 | 1.74 | 0.41 | D | 0 | 0 |
| | GLU190 | OE2 | 40 | 0.87 | 2.06 | 1.14 | A | 1 | 0 |
| | GLY135 | O | 40 | 5.22 | 6.33 | 0.66 | A | 0 | 1 |
| | SER136 | OG | 40 | 3.41 | 4.15 | 0.66 | D | 0 | 0 |
| | TYR98 | OH | 40 | 1.26 | 3.18 | 1.70 | A | 0 | 0 |

| | | | | | | | | | |
|---|---|---|---|---|---|---|---|---|---|
| **1HWI** | ARG590 | NH2 | 40 | 6.31 | 8.49 | 1.30 | D | 0 | 0 |
| | ASN755 | ND2 | 40 | 9.43 | 9.67 | 0.17 | D | 0 | 1 |
| | ASP690 | OD2 | 40 | 9.96 | 12.24 | 1.49 | A | 0 | 0 |
| | GLU559 | OE2 | 40 | 5.51 | 6.36 | 0.68 | A | 0 | 0 |
| | LYS691 | NZ | 40 | 6.78 | 8.20 | 1.39 | D | 0 | 0 |
| | LYS692 | NZ | 40 | 7.02 | 7.20 | 0.16 | D | 1 | 0 |
| | LYS735 | NZ | 40 | 10.01 | 11.77 | 1.18 | D | 1 | 0 |
| | SER684 | OG | 40 | 5.63 | 6.86 | 0.86 | D | 0 | 0 |
| **1IVB** | ARG116 | NH2 | 40 | 3.83 | 5.37 | 0.95 | D | 1 | 0 |
| | ARG292 | NH2 | 40 | 5.79 | 6.33 | 0.38 | D | 1 | 0 |
| | ARG374 | NH1 | 40 | 12.68 | 13.07 | 0.55 | D | 1 | 0 |
| | ARG374 | NH2 | 40 | 6.36 | 7.82 | 1.38 | D | 1 | 0 |
| | GLU117 | OE2 | 40 | 2.04 | 2.89 | 0.96 | A | 0 | 0 |
| | TYR409 | OH | 40 | 12.10 | 13.12 | 0.61 | D | 0 | 0 |
| **1IVD** | ARG118 | NH1 | 40 | 1.52 | 2.47 | 0.55 | D | 1 | 0 |
| | ARG118 | NH2 | 40 | 5.28 | 5.55 | 0.19 | D | 1 | 0 |
| | ARG292 | NH2 | 40 | 7.35 | 7.78 | 0.32 | D | 1 | 0 |
| **1IVE** | ARG118 | NH2 | 18 | 0.08 | 0.13 | 0.04 | D | 1 | 0 |
| | ARG152 | NH1 | 40 | 5.30 | 5.87 | 0.37 | D | 0 | 0 |
| | ARG292 | NH2 | 40 | 3.91 | 4.48 | 0.59 | D | 1 | 0 |
| | ARG371 | NH1 | 40 | 1.72 | 2.56 | 0.57 | D | 1 | 0 |
| | ARG371 | NH2 | 40 | 2.16 | 3.14 | 0.60 | D | 1 | 0 |
| **1IVF** | ARG118 | NH1 | 40 | 6.88 | 7.54 | 0.50 | D | 1 | 0 |
| | ARG152 | NE | 40 | 1.32 | 2.79 | 0.90 | D | 0 | 0 |
| | ARG152 | NH2 | 40 | 0.51 | 0.91 | 0.33 | D | 0 | 0 |
| | ARG224 | NE | 40 | 2.18 | 3.64 | 0.91 | D | 0 | 0 |
| | ARG371 | NH1 | 40 | 4.94 | 5.75 | 0.77 | D | 1 | 0 |
| | ARG371 | NH2 | 40 | 8.14 | 9.39 | 1.29 | D | 1 | 0 |
| | ASP151 | OD1 | 40 | 2.60 | 2.74 | 0.11 | A | 0 | 0 |
| | GLU119 | OE2 | 40 | 3.77 | 5.66 | 1.51 | A | 0 | 0 |
| | GLU276 | OE1 | 40 | 5.92 | 7.28 | 1.02 | A | 0 | 0 |
| **1JLA** | LYS101 | O | 40 | 2.07 | 3.01 | 0.62 | A | 0 | 1 |
| **1K1J** | ASP189 | OD1 | 40 | 16.49 | 16.62 | 0.10 | A | 1 | 0 |
| | ASP189 | OD2 | 40 | 15.76 | 15.82 | 0.06 | A | 1 | 0 |
| | GLY216 | N | 40 | 11.28 | 11.97 | 0.45 | D | 0 | 1 |
| | GLY216 | O | 40 | 9.87 | 10.34 | 0.29 | A | 0 | 1 |
| | GLY219 | N | 40 | 7.69 | 8.38 | 0.55 | D | 0 | 1 |
| | GLY219 | O | 40 | 15.52 | 16.48 | 0.56 | A | 0 | 0 |
| **1K3U** | ASP60 | OD2 | 40 | 2.05 | 2.69 | 0.81 | A | 0 | 0 |
| | GLY184 | N | 40 | 4.22 | 5.07 | 0.86 | D | 0 | 0 |
| | GLY213 | N | 40 | 2.91 | 5.47 | 1.81 | D | 0 | 0 |
| | GLY234 | N | 40 | 11.76 | 11.94 | 0.11 | D | 0 | 0 |

| | | | | | | | | | |
|---|---|---|---|---|---|---|---|---|---|
| | ILE214 | N | 40 | 1.56 | 3.95 | 1.41 | D | 0 | 0 |
| | PHE212 | N | 40 | 0.26 | 0.52 | 0.17 | D | 0 | 0 |
| | SER235 | N | 40 | 2.57 | 3.39 | 1.01 | D | 0 | 0 |
| | SER235 | OG | 40 | 9.69 | 10.44 | 0.77 | D | 0 | 0 |
| | TYR175 | OH | 40 | 17.84 | 19.52 | 1.41 | D | 0 | 1 |
| **1KE5** | ASP86 | N | 40 | 0.43 | 0.68 | 0.14 | D | 0 | 1 |
| | GLU81 | O | 40 | 7.72 | 8.14 | 0.37 | A | 0 | 1 |
| | LEU83 | N | 40 | 7.93 | 8.21 | 0.28 | D | 0 | 1 |
| **1L2S** | ALA318 | N | 40 | 9.21 | 10.04 | 0.68 | D | 0 | 0 |
| | ALA318 | O | 40 | 4.83 | 5.25 | 0.44 | A | 0 | 1 |
| | ASN152 | ND2 | 40 | 0.98 | 1.79 | 0.96 | D | 0 | 1 |
| | SER64 | N | 40 | 10.55 | 11.86 | 0.84 | D | 0 | 0 |
| | SER64 | OG | 40 | 10.41 | 11.72 | 1.17 | D | 0 | 1 |
| **1L7F** | ARG118 | NH1 | 40 | 3.24 | 3.96 | 0.49 | D | 1 | 0 |
| | ARG118 | NH2 | 40 | 4.35 | 6.79 | 1.56 | D | 1 | 0 |
| | ARG152 | NH2 | 40 | 6.45 | 7.32 | 0.82 | D | 0 | 0 |
| | ARG292 | NH1 | 40 | 6.25 | 7.55 | 0.76 | D | 1 | 0 |
| | ARG292 | NH2 | 40 | 8.92 | 9.21 | 0.42 | D | 1 | 0 |
| | ARG371 | NH1 | 40 | 10.28 | 12.82 | 1.62 | D | 1 | 0 |
| | ARG371 | NH2 | 40 | 7.05 | 9.44 | 1.66 | D | 1 | 0 |
| | ASP151 | O | 40 | 2.65 | 3.00 | 0.35 | A | 0 | 0 |
| | ASP151 | OD1 | 40 | 0.79 | 1.14 | 0.24 | A | 1 | 0 |
| | GLU227 | OE1 | 40 | 10.20 | 10.34 | 0.11 | A | 1 | 0 |
| | TRP178 | O | 40 | 9.54 | 9.83 | 0.29 | A | 0 | 0 |
| **1LPZ** | ASP189 | OD1 | 40 | 10.62 | 11.32 | 0.86 | A | 1 | 0 |
| | ASP189 | OD2 | 40 | 13.27 | 13.89 | 0.39 | A | 1 | 0 |
| | GLY219 | O | 40 | 13.15 | 14.22 | 0.90 | A | 0 | 0 |
| **1LQD** | ASP189 | OD1 | 40 | 5.58 | 7.64 | 1.21 | A | 1 | 0 |
| | ASP189 | OD2 | 40 | 12.33 | 13.50 | 1.16 | A | 1 | 0 |
| | GLY219 | O | 40 | 6.97 | 7.39 | 0.38 | A | 0 | 0 |
| **1M2Z** | ARG611 | NH2 | 40 | 7.99 | 8.26 | 0.28 | D | 0 | 0 |
| | ASN564 | ND2 | 40 | 3.41 | 3.68 | 0.25 | D | 0 | 1 |
| | ASN564 | OD1 | 40 | 6.13 | 6.76 | 0.73 | A | 0 | 1 |
| | GLN570 | NE2 | 40 | 6.80 | 6.99 | 0.13 | D | 0 | 1 |
| | GLN642 | OE1 | 40 | 10.60 | 11.54 | 0.82 | A | 0 | 1 |
| | THR739 | OG1 | 4 | 0.06 | 0.38 | 0.22 | A | 0 | 1 |
| **1ML1** | ASN11 | ND2 | 40 | 11.02 | 11.42 | 0.34 | D | 0 | 0 |
| | GLH167 | OE2 | 40 | 11.62 | 12.17 | 0.56 | A | 0 | 0 |
| | GLY173 | N | 40 | 11.65 | 13.22 | 1.51 | D | 0 | 1 |
| | GLY234 | N | 40 | 11.43 | 12.65 | 0.79 | D | 0 | 0 |
| | GLY235 | N | 40 | 7.04 | 9.67 | 1.74 | D | 0 | 0 |
| | HIP95 | NE2 | 40 | 25.67 | 26.54 | 0.60 | D | 1 | 0 |

| | | | | | | | | | |
|---|---|---|---|---|---|---|---|---|---|
| | LYS13 | NZ | 40 | 14.03 | 15.06 | 0.79 | D | 1 | 0 |
| | SER213 | N | 40 | 7.53 | 8.80 | 0.80 | D | 0 | 1 |
| **1MQ6** | GLY216 | N | 40 | 6.11 | 6.14 | 0.03 | D | 0 | 1 |
| | GLY218 | N | 40 | 3.70 | 3.83 | 0.15 | D | 0 | 1 |
| **1MTS** | ASP189 | OD1 | 40 | 13.48 | 14.74 | 0.77 | A | 1 | 0 |
| | ASP189 | OD2 | 40 | 20.17 | 21.17 | 0.65 | A | 1 | 0 |
| | GLY219 | O | 40 | 8.94 | 9.38 | 0.49 | A | 0 | 0 |
| **1N2J** | GLN164 | NE2 | 40 | 3.31 | 3.49 | 0.14 | D | 0 | 0 |
| | GLN164 | OE1 | 40 | 10.22 | 11.20 | 0.79 | A | 0 | 1 |
| | GLN72 | NE2 | 40 | 7.42 | 8.67 | 0.81 | D | 0 | 1 |
| | GLN72 | OE1 | 40 | 7.67 | 8.56 | 0.66 | A | 0 | 1 |
| **1N2V** | ASP156 | OD1 | 40 | 6.39 | 7.80 | 1.37 | A | 0 | 0 |
| | ASP156 | OD2 | 40 | 9.20 | 10.72 | 0.88 | A | 0 | 0 |
| | GLN203 | NE2 | 40 | 5.88 | 6.53 | 0.44 | D | 0 | 1 |
| | GLY230 | N | 40 | 12.36 | 12.74 | 0.35 | D | 0 | 1 |
| **1N46** | ARG316 | NH1 | 40 | 14.78 | 15.07 | 0.21 | D | 0 | 0 |
| | ARG320 | NH1 | 40 | 19.93 | 21.74 | 1.39 | D | 0 | 0 |
| | ARG320 | NH2 | 40 | 21.58 | 22.40 | 0.48 | D | 1 | 0 |
| | ASN331 | N | 40 | 12.00 | 13.25 | 1.25 | D | 0 | 1 |
| | HID435 | NE2 | 40 | 11.88 | 12.23 | 0.26 | A | 0 | 1 |
| **1OF1** | GLN125 | NE2 | 40 | 12.32 | 12.94 | 0.54 | D | 0 | 1 |
| | GLN125 | OE1 | 40 | 9.92 | 10.72 | 0.62 | A | 0 | 1 |
| | GLU225 | OE1 | 40 | 6.92 | 7.19 | 0.33 | A | 0 | 0 |
| | TYR101 | OH | 40 | 6.24 | 7.00 | 0.56 | D | 0 | 1 |
| **1OWE** | ASP205 | OD1 | 40 | 16.26 | 18.08 | 1.18 | A | 1 | 0 |
| | ASP205 | OD2 | 40 | 17.84 | 19.21 | 0.92 | A | 1 | 0 |
| | GLY234 | O | 40 | 9.63 | 9.85 | 0.18 | A | 0 | 0 |
| | SER206 | OG | 40 | 9.59 | 9.74 | 0.15 | A | 0 | 0 |
| **1OYT** | ASP189 | OD1 | 40 | 9.02 | 10.67 | 1.92 | A | 1 | 0 |
| | ASP189 | OD2 | 40 | 10.53 | 11.69 | 0.72 | A | 1 | 0 |
| | GLY216 | N | 40 | 8.41 | 9.33 | 0.58 | D | 0 | 1 |
| | GLY219 | O | 40 | 6.85 | 8.72 | 1.28 | A | 0 | 0 |
| **1PMN** | MET149 | N | 40 | 7.96 | 9.00 | 0.71 | D | 0 | 1 |
| | MET149 | O | 40 | 6.75 | 7.64 | 0.70 | A | 0 | 1 |
| **1Q1G** | ASP206 | OD1 | 40 | 8.38 | 10.38 | 1.25 | A | 0 | 0 |
| | GLU184 | OE1 | 40 | 3.03 | 7.51 | 2.66 | A | 0 | 0 |
| | GLU184 | OE2 | 40 | 5.89 | 7.03 | 0.88 | A | 0 | 0 |
| | MET183 | N | 40 | 0.89 | 1.41 | 0.40 | D | 0 | 1 |
| | SER91 | OG | 40 | 0.77 | 1.22 | 0.27 | A | 0 | 0 |
| **1Q41** | ASP133 | O | 40 | 14.34 | 15.01 | 0.52 | A | 0 | 1 |
| | VAL135 | N | 40 | 14.28 | 15.93 | 0.96 | D | 0 | 1 |
| | VAL135 | O | 40 | 8.38 | 9.77 | 0.83 | A | 0 | 1 |

| | | | | | | | | | |
|---|---|---|---|---|---|---|---|---|---|
| **1QHI** | ARG176 | NE | 40 | 4.88 | 8.31 | 3.41 | D | 0 | 0 |
| | ARG176 | NH2 | 40 | 0.12 | 4.44 | 2.58 | D | 0 | 0 |
| | GLN125 | NE2 | 40 | 2.46 | 3.88 | 1.78 | D | 0 | 1 |
| | GLN125 | OE1 | 40 | 12.89 | 13.29 | 0.35 | A | 0 | 1 |
| | GLU83 | OE2 | 40 | 3.28 | 4.03 | 0.60 | A | 0 | 0 |
| **1ROB** | GLN11 | NE2 | 40 | 1.30 | 1.84 | 0.62 | D | 0 | 1 |
| | HIP119 | ND1 | 40 | 1.79 | 2.05 | 0.25 | D | 1 | 0 |
| | HIP12 | NE2 | 40 | 2.54 | 3.24 | 0.56 | D | 1 | 0 |
| | LYS41 | NZ | 40 | 1.52 | 3.21 | 1.56 | D | 0 | 0 |
| | PHE120 | N | 40 | 2.01 | 3.02 | 0.98 | D | 0 | 0 |
| | THR45 | N | 40 | 5.69 | 7.75 | 2.39 | D | 0 | 1 |
| | THR45 | OG1 | 40 | 6.62 | 7.54 | 0.94 | D | 0 | 1 |
| **1S19** | ARG274 | NH1 | 40 | 8.04 | 9.69 | 0.97 | D | 0 | 0 |
| | HID305 | NE2 | 40 | 7.29 | 7.60 | 0.21 | A | 0 | 1 |
| | HIE397 | NE2 | 40 | 5.15 | 5.68 | 0.61 | D | 0 | 1 |
| | SER237 | OG | 40 | 9.55 | 10.53 | 0.57 | A | 0 | 1 |
| | SER278 | OG | 40 | 7.59 | 8.46 | 0.58 | A | 0 | 1 |
| | TYR143 | OH | 40 | 6.56 | 7.02 | 0.38 | D | 0 | 1 |
| **1TOW** | ARG126 | NE | 40 | 3.46 | 6.38 | 1.92 | D | 1 | 0 |
| | ARG126 | NH2 | 40 | 5.60 | 6.81 | 1.44 | D | 1 | 0 |
| | TYR128 | OH | 40 | 4.98 | 6.46 | 0.93 | D | 0 | 0 |
| **1TT1** | ALA142 | N | 40 | 13.08 | 13.58 | 0.33 | D | 0 | 0 |
| | ALA91 | N | 40 | 16.28 | 16.57 | 0.21 | D | 0 | 0 |
| | ARG96 | NH1 | 40 | 27.36 | 27.92 | 0.53 | D | 1 | 0 |
| | ARG96 | NH2 | 40 | 13.94 | 17.94 | 2.40 | D | 1 | 0 |
| | GLU191 | OE1 | 40 | 20.28 | 21.47 | 1.19 | A | 1 | 0 |
| | PRO89 | O | 40 | 16.71 | 18.34 | 1.10 | A | 0 | 0 |
| | THR143 | N | 40 | 9.22 | 11.25 | 1.18 | D | 0 | 0 |
| | THR143 | OG1 | 40 | 10.14 | 11.64 | 1.34 | D | 0 | 0 |
| **1U4D** | ALA208 | N | 40 | 8.00 | 8.60 | 0.48 | D | 0 | 1 |
| | ASP270 | OD1 | 40 | 0.04 | 0.16 | 0.08 | A | 0 | 0 |
| | GLU206 | O | 40 | 3.72 | 4.02 | 0.23 | A | 0 | 1 |
| | LYS158 | NZ | 40 | 2.32 | 3.28 | 0.65 | D | 0 | 0 |
| **1UKZ** | ARG107 | NH1 | 40 | 15.19 | 18.45 | 1.89 | D | 0 | 0 |
| | ARG107 | NH1 | 40 | 10.11 | 13.53 | 2.62 | D | 1 | 0 |
| | ARG107 | NH2 | 40 | 8.62 | 9.01 | 0.27 | D | 1 | 0 |
| | ARG148 | NH2 | 40 | 4.41 | 5.12 | 0.51 | D | 1 | 0 |
| | ARG52 | NH1 | 40 | 4.96 | 5.39 | 0.70 | D | 0 | 0 |
| | ARG52 | NH2 | 40 | 4.27 | 4.76 | 0.32 | D | 0 | 0 |
| | GLN111 | NE2 | 40 | 8.79 | 9.37 | 0.48 | D | 0 | 1 |
| | GLN111 | OE1 | 40 | 7.98 | 12.06 | 2.51 | A | 0 | 1 |
| | GLN74 | O | 40 | 13.54 | 13.77 | 0.20 | A | 0 | 1 |

| | | | | | | | | | |
|---|---|---|---|---|---|---|---|---|---|
| | GLY104 | O | 40 | 15.28 | 15.70 | 0.26 | A | 0 | 1 |
| | VAL76 | N | 40 | 17.80 | 20.13 | 1.87 | D | 0 | 1 |
| **1ULB** | ASN243 | ND2 | 40 | 2.67 | 2.88 | 0.13 | D | 0 | 1 |
| | ASN243 | ND2 | 40 | 4.42 | 5.01 | 0.39 | D | 0 | 1 |
| | GLU201 | OE1 | 40 | 7.33 | 13.09 | 3.52 | A | 0 | 0 |
| | GLU201 | OE2 | 40 | 10.82 | 11.86 | 0.71 | A | 0 | 0 |
| | LYS244 | NZ | 40 | 11.72 | 12.42 | 0.57 | D | 0 | 0 |
| **1UNL** | CYS83 | N | 40 | 7.11 | 7.28 | 0.19 | D | 0 | 1 |
| | CYS83 | O | 40 | 4.26 | 5.44 | 0.78 | A | 0 | 1 |
| | GLN130 | O | 40 | 0.19 | 1.15 | 0.74 | A | 0 | 1 |
| **1UOU** | ARG202 | NH1 | 40 | 9.12 | 11.15 | 1.22 | D | 0 | 0 |
| | ARG202 | NH2 | 40 | 9.88 | 12.07 | 1.48 | D | 0 | 0 |
| | HID116 | NE2 | 40 | 12.04 | 13.41 | 0.81 | A | 0 | 1 |
| | LYS221 | NZ | 40 | 17.51 | 18.22 | 0.68 | D | 0 | 0 |
| | SER117 | O | 40 | 1.08 | 1.56 | 0.30 | A | 0 | 0 |
| | SER217 | OG | 40 | 15.85 | 17.74 | 1.16 | A | 0 | 1 |
| **1V0P** | ASP85 | OD1 | 40 | 0.03 | 0.50 | 0.77 | A | 0 | 0 |
| | GLN129 | O | 40 | 4.04 | 4.94 | 0.72 | A | 0 | 1 |
| | LEU82 | N | 40 | 3.12 | 4.36 | 0.79 | D | 0 | 1 |
| | LEU82 | O | 40 | 2.68 | 3.35 | 0.53 | A | 0 | 1 |
| **1W1P** | TRP97 | N | 40 | 1.25 | 2.32 | 0.69 | D | 0 | 1 |
| | TYR214 | OH | 40 | 3.00 | 3.41 | 0.35 | D | 0 | 1 |
| **1W2G** | ARG74 | NH1 | 40 | 15.40 | 15.62 | 0.17 | D | 0 | 0 |
| | ARG74 | NH2 | 40 | 11.50 | 11.93 | 0.30 | D | 0 | 0 |
| | ASN100 | OD1 | 40 | 6.76 | 6.90 | 0.10 | A | 0 | 1 |
| **1X8X** | ASP182 | OD1 | 40 | 21.43 | 22.02 | 0.39 | A | 0 | 0 |
| | ASP81 | OD1 | 40 | 5.35 | 5.57 | 0.16 | A | 1 | 0 |
| | GLN179 | OE1 | 40 | 15.38 | 15.79 | 0.27 | A | 0 | 0 |
| | TYR175 | OH | 40 | 14.57 | 15.86 | 0.75 | A | 0 | 0 |
| | TYR37 | OH | 40 | 9.31 | 10.45 | 0.88 | D | 0 | 1 |
| **1YDR** | VAL123 | N | 40 | 7.78 | 8.34 | 0.33 | D | 0 | 1 |
| **1YDS** | ASP184 | OD2 | 40 | 1.04 | 2.09 | 1.02 | A | 1 | 0 |
| | GLU170 | O | 40 | 5.22 | 5.86 | 0.49 | A | 0 | 0 |
| | VAL123 | N | 40 | 4.55 | 5.31 | 0.49 | D | 0 | 1 |
| **1YDT** | ASN171 | OD1 | 40 | 2.21 | 2.44 | 0.14 | A | 0 | 0 |
| | GLU170 | O | 40 | 1.03 | 2.20 | 0.80 | A | 0 | 0 |
| | VAL123 | N | 40 | 5.51 | 5.85 | 0.25 | D | 0 | 1 |
| **1YV3** | GLY240 | N | 40 | 17.24 | 20.64 | 2.01 | D | 0 | 1 |
| | LEU262 | O | 40 | 24.42 | 26.41 | 1.18 | A | 0 | 1 |
| | SER456 | N | 40 | 6.20 | 6.36 | 0.16 | D | 0 | 1 |
| **1YVF** | TYR448 | N | 40 | 3.28 | 3.70 | 0.37 | D | 0 | 1 |
| **1YWR** | ASP168 | OD2 | 40 | 1.44 | 1.58 | 0.11 | A | 1 | 0 |

| | | | | | | | | | |
|---|---|---|---|---|---|---|---|---|---|
| | LYS53 | NZ | 40 | 2.82 | 3.85 | 0.64 | D | 0 | 0 |
| | MET109 | N | 40 | 8.47 | 9.33 | 0.60 | D | 0 | 1 |
| | MET109 | O | 40 | 3.89 | 5.01 | 0.73 | A | 0 | 1 |
| **2ACK** | HIP440 | NE2 | 6 | 0.35 | 2.11 | 1.29 | D | 0 | 1 |
| | SER200 | OG | 40 | 0.47 | 1.46 | 0.65 | A | 0 | 1 |
| **2BR1** | CYS87 | N | 40 | 5.25 | 6.52 | 0.78 | D | 0 | 1 |
| **2MCP** | ARG52 | NH1 | 40 | 0.52 | 0.71 | 0.15 | D | 1 | 0 |
| | TYR33 | OH | 20 | 0.33 | 1.25 | 0.54 | D | 0 | 0 |
| **2PCP** | TRP97 | O | 40 | 13.28 | 14.07 | 0.62 | A | 0 | 0 |
| **3PTB** | ASP189 | OD1 | 40 | 13.26 | 13.73 | 0.28 | A | 1 | 0 |
| | ASP189 | OD2 | 40 | 14.60 | 15.09 | 0.53 | A | 1 | 0 |
| | GLY219 | O | 40 | 9.01 | 10.19 | 0.77 | A | 0 | 0 |
| **4TS1** | ASP176 | OD1 | 40 | 20.98 | 21.81 | 0.55 | A | 0 | 0 |
| | ASP78 | OD1 | 40 | 8.78 | 9.98 | 0.70 | A | 1 | 0 |
| | GLN173 | OE1 | 40 | 10.09 | 10.73 | 0.68 | A | 0 | 0 |
| | TYR169 | OH | 40 | 15.28 | 15.89 | 0.35 | A | 0 | 0 |
| | TYR34 | OH | 40 | 1.55 | 1.99 | 0.69 | D | 0 | 1 |

***Supplementary Figure 3*** *Structures of protein-ligand complexes that do not poses any robust hydrogen bond (all the bond are weaker then 4 kcal/mole). Weak hydrogen bonds ($W_{QB}$ < 4 kcal/mol) marked in green. The first group (green, PDB codes: 1JLA and 1C1B) represents complexes of reverse transcriptase. Activity data of molecules: IC50 = 6 nM for 1JLA[11] and EC50 = 0.6 nM for 1C1B[12]. The second group (orange, PDB codes: 1HGH_2 and 1W1P) represent ligands with low activity. 1W1P has IC50 value of 5 mM[13]. 1HGH_2 is a ligand bound in the secondary binding side of hemagglutinin with considerably lower affinity that the primary binding site.[2] Third group (violet, PDB codes: 2ACK and 2MCP) represents complexes stabilized by cation-pi interaction, making hydrogen bonds weak. The interaction between cation and the centroid of tryptophan was marked in yellow. The last group (magenta, PDB codes: 1F0S, 1G9V and 1YVF) represents complexes stabilized by water mediated hydrogen bonds. The water network is marked as red '+' and the water mediated interactions are marked as yellow. Activity data of molecules $K_i$ = 18 nM for 1F0S[14] and IC50 = 100 nM for 1YVF.[15]*

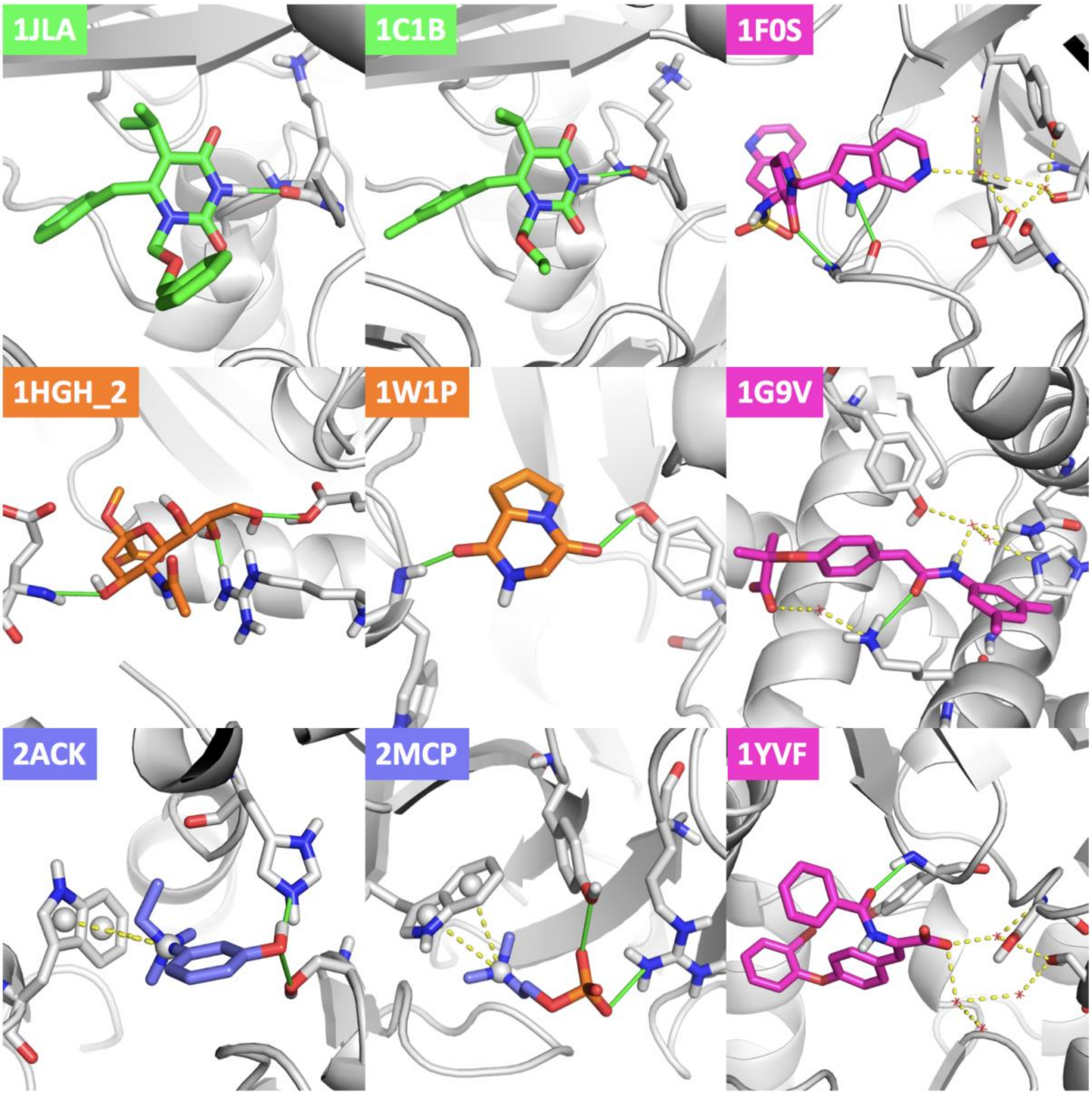

***Supplementary Table 3** Classification of structures in Iridium DS based on function*

| Category | PDB codes |
|---|---|
| Enzymes | 1ai5, 1b9v, 1cvu, 1dds, 1ezq, 1f0s, 1f0t, 1f0u, 1fh8, 1fh9, 1fhd, 1fvt, 1gm8, 1h1p, 1h1s, 1hwi, 1ivb, 1ivd, 1ive, 1ivf, 1k1j, 1ke5, 1l2s, 1l7f, 1lpz, 1lqd, 1ml1, 1mq6, 1mts, 1n2j, 1n2v, 1of1, 1owe, 1oyt, 1pmn, 1q1g, 1q41, 1qhi, 1rob, 1u4d, 1ukz, 1ulb, 1unl, 1uou, 1v0p, 1w1p, 1w2g, 1x8x, 1ydr, 1yds, 1ydt, 1ywr, 2ack, 2br1, 3ptb, 4ts1 |
| Kinases | 1fvt, 1h1p, 1h1s, 1ke5, 1of1, 1pmn, 1q41, 1qhi, 1u4d, 1ukz, 1w2g, 1ydr, 1yds, 1ydt, 1ywr, 2br1 |
| Proteases | 1ezq, 1f0s, 1f0t, 1f0u, 1k1j, 1lpz, 1lqd, 1mq6, 1mts, 3ptb |
| Receptors | Nuclear receptors: 1a28, 1exa, 1fcx, 1fcz, 1fm6, 1m2z, 1n46, 1s19<br>Glutamate receptor: 1tt1 |
| Carbohydrate binding prot. | 1b9v, 1br6, 1fh8, 1fh9, 1fhd, 1hgh_1, 1hgi, 1hgj, 1ivb, 1ivd, 1ive, 1ivf, 1l7f, 1w1p |
| Allosteric sites | 1c1b, 1g9v, 1hgh_2, 1jla, 1k3u, 1m2z, 1yv3, 1yvf |

***Supplementary Table 4** Two sample Kolmogorov-Smirnov statistical test of distribution of WQB values across the four types of binding sites defined in Figure 1.*

| **p-values** | Enzymes | Nuclear receptors | Carbohydrate binding prot. | Allosteric sites |
|---|---|---|---|---|
| Enzymes | | | | |
| Nuclear receptors | 0.071 | | | |
| Carbohydrate binding prot. | 0.055 | 3.3e-3 | | |
| Allosteric sites | 0.011 | 8.2e-4 | 0.24 | |

***Supplementary Table 5** The average number of total and robust hydrogen bonds for the four types of binding sites defined in Figure 1.*

| **Avg. HBs** | Total | Robust |
|---|---|---|
| Enzymes | 4.52 | 2.73 |
| Nuclear receptors | 3.86 | 3 |
| Carbohydrate binding prot. | 6.79 | 3.14 |
| Allosteric sites | 3.13 | 1.25 |

***Supplementary Figure 4** Distribution of relative distance between the center of mass (CM) of all atoms and: the center of all heteroatoms (N and O) in blue, the center of atoms making hydrogen bonds in green, the center of atoms making only strong and medium interactions in pink, for all of calculated complexes. Above the plot, an example of depicted molecule with marked corresponding vectors, is presented. Relative distance calculated as a result of division of the length of vector span between the CM and the center of selected atoms, and the length of vector span between the CM and the furthest atom in the structure.*

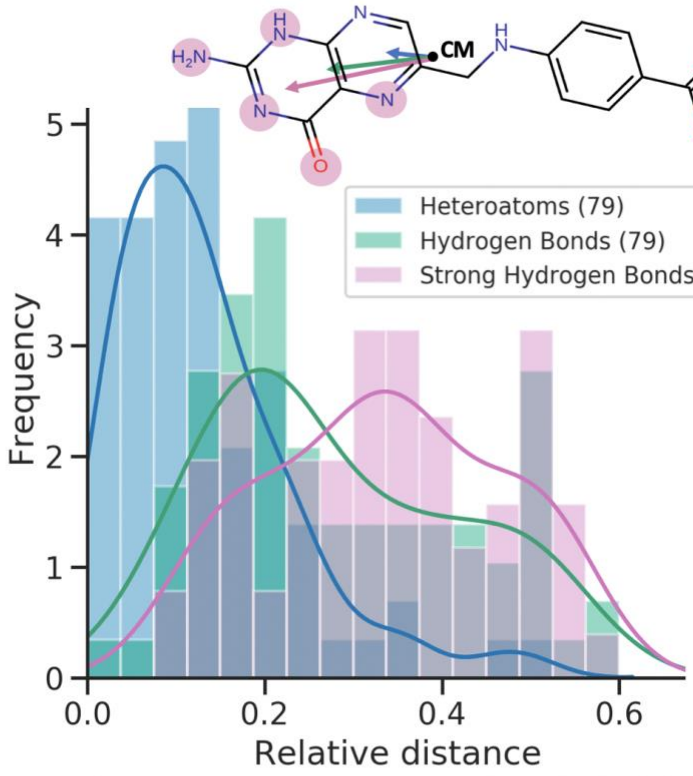

***Supplementary Figure 5*** *Depicted ligands from Iridium DS, with corresponding PDB codes. Atoms that make hydrogen bonds with the protein were highlighted and the* $W_{QB}$ *value associated with that bond is written below the atom (multiple values in cases when the atom is making more than one hydrogen bond). The colour of highlight symbolizes fragment-sized group of atoms. Atom were grouped based on the distance with the cut-off of 4 Å and a few classifications were corrected based on visual inspection.*

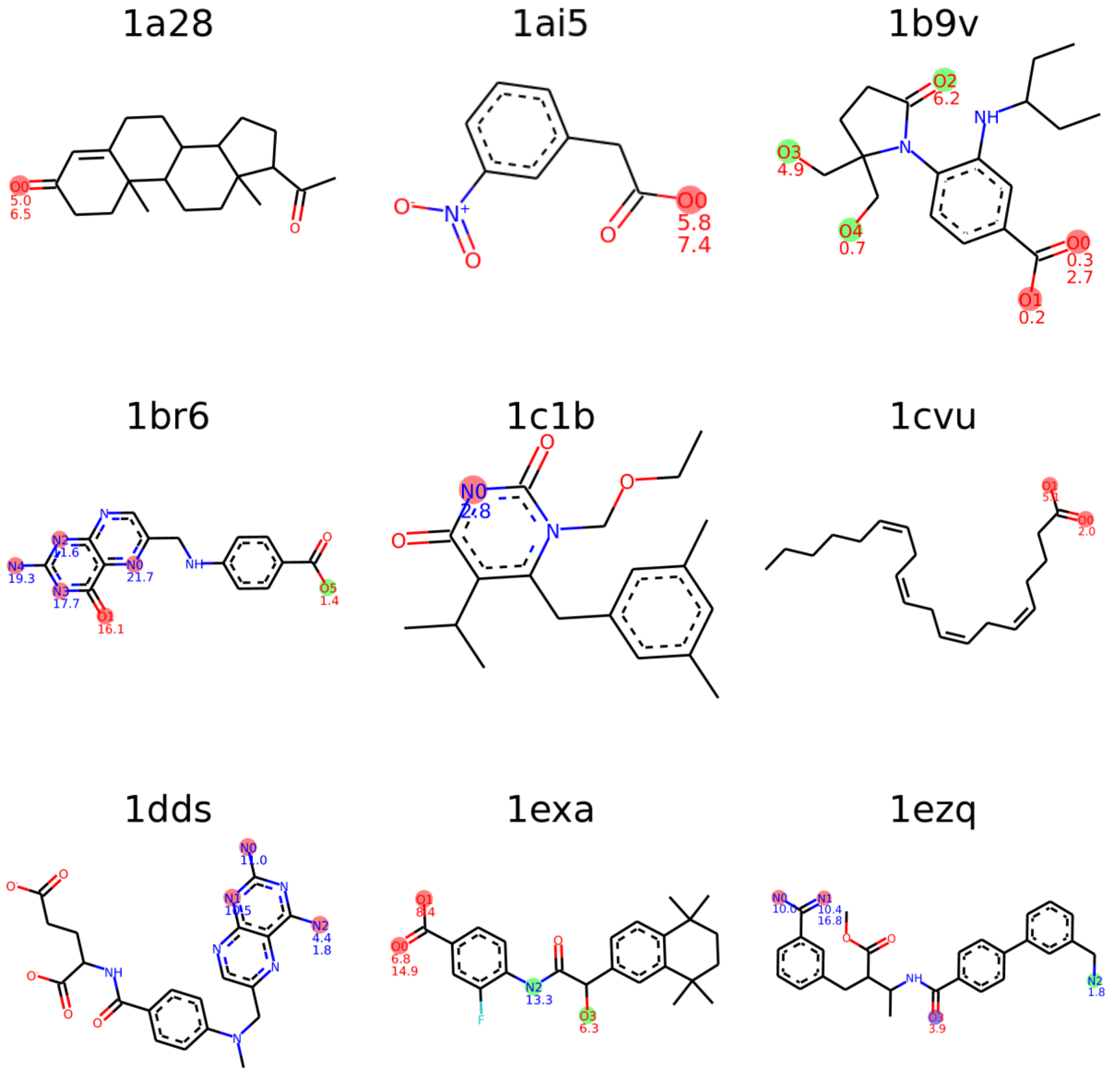

1f0s

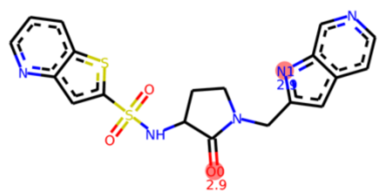

1f0t

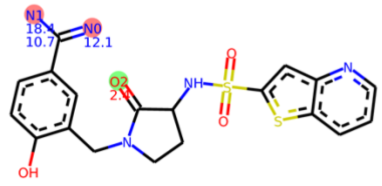

1f0u

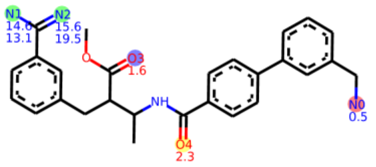

1fcx

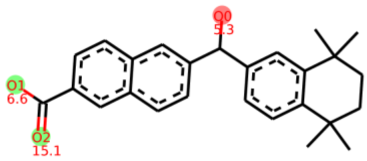

1fcz

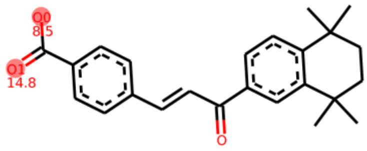

1fh8

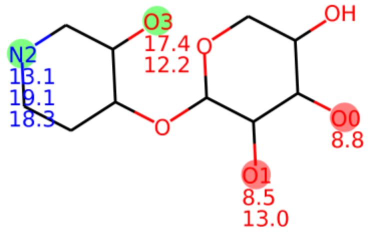

1fh9

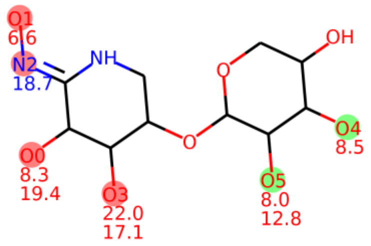

1fhd

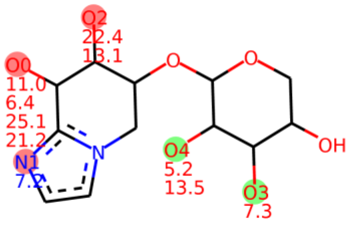

1fm6

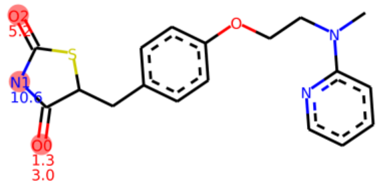

1fvt

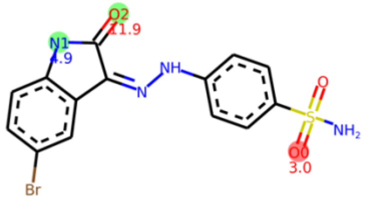

1g9v

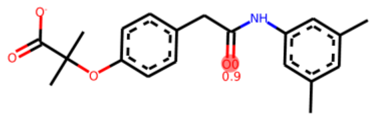

1gm8

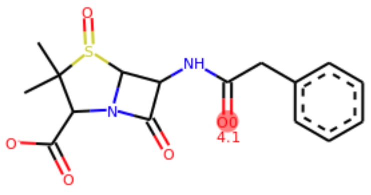

1h1p

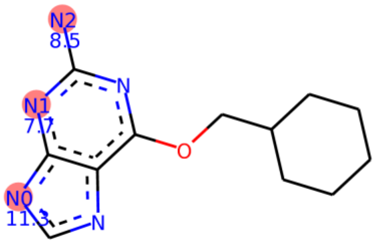


1h1s

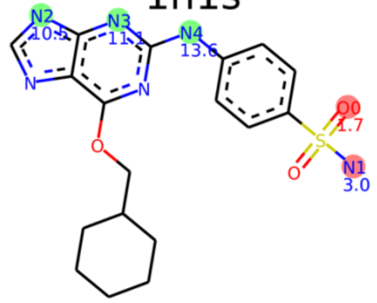


1hgh_1

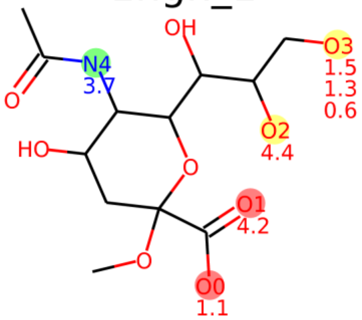


1hgh_2

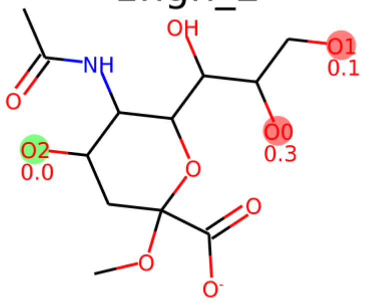


1hgi

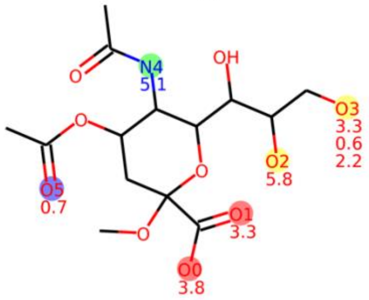


1hgj

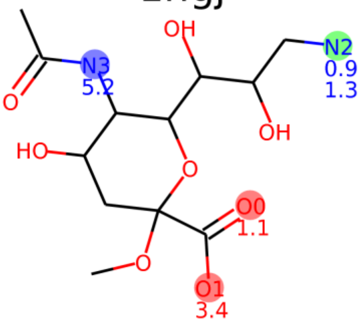


1hwi

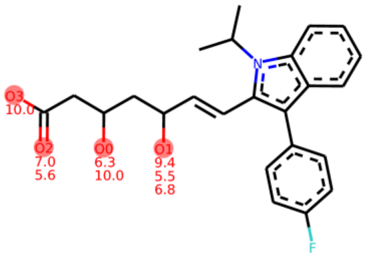


1ivb

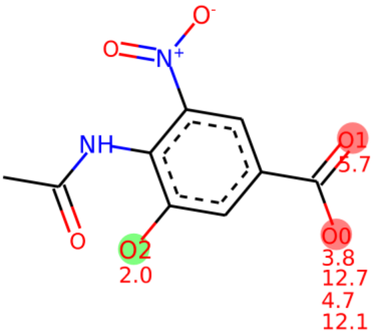


1ivd

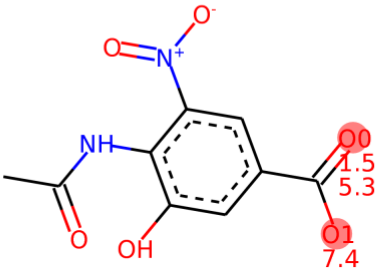


1ive

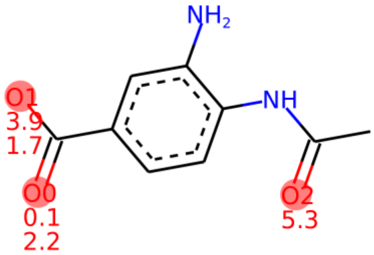


1ivf

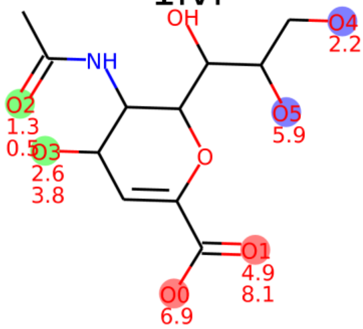


1jla

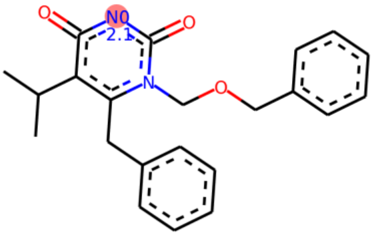

1k1j

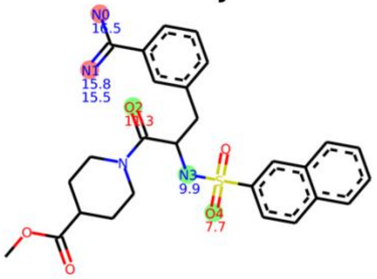

1k3u

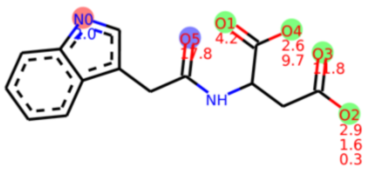

1ke5

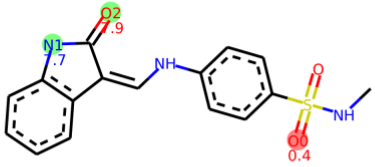

1l2s

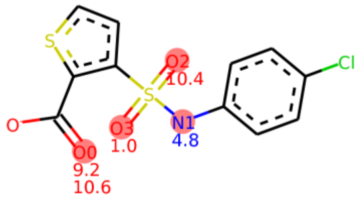

1l7f

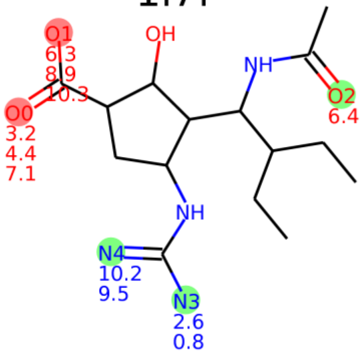

1lpz

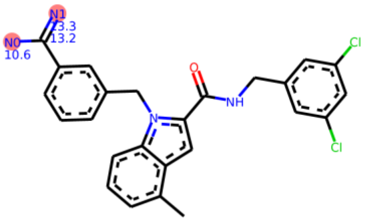

1lqd

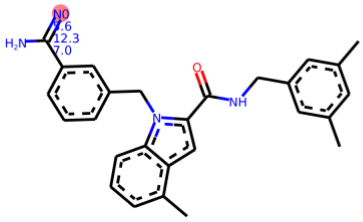

1m2z

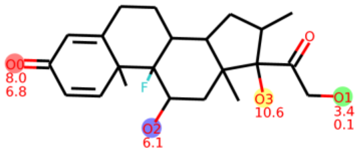

1ml1

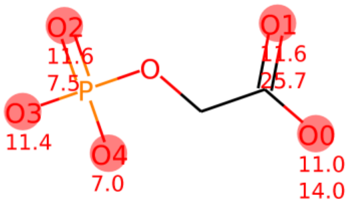

1mq6

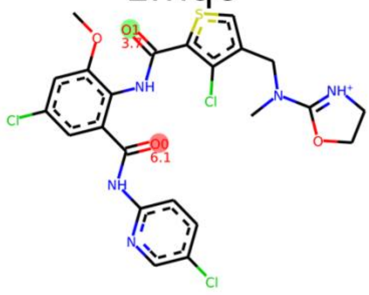

1mts

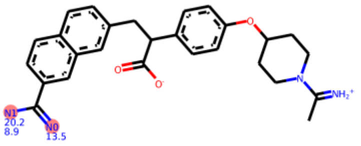

1n2j

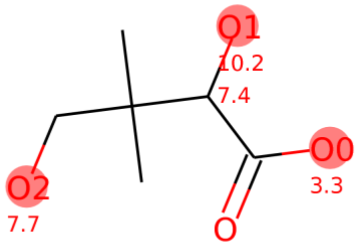

1n2v

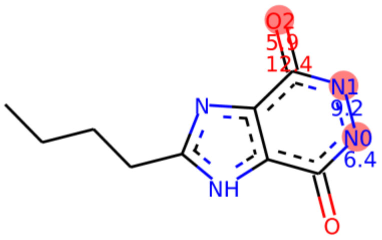

1n46

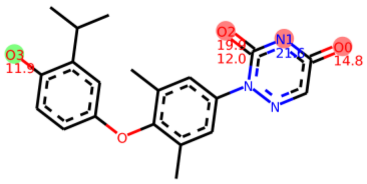

1of1

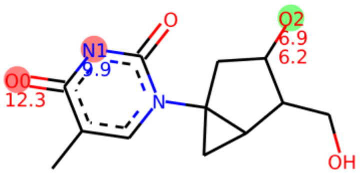

1owe

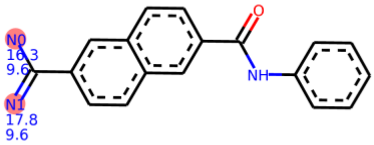

1oyt

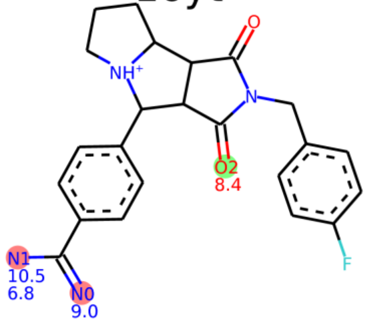

1pmn

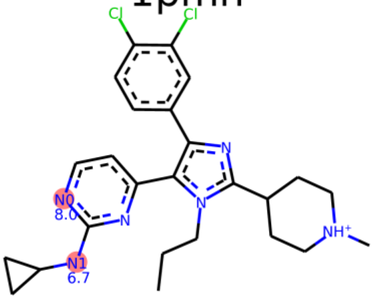

1q1g

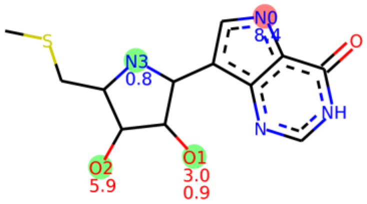

1q41

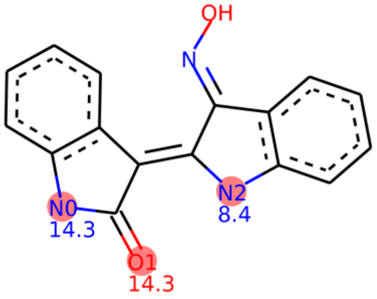

1qhi

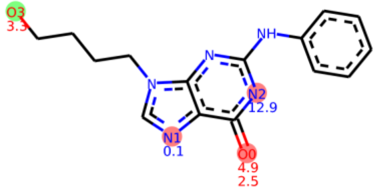

1rob

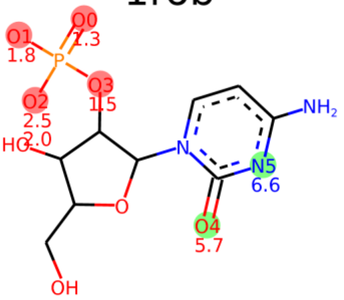

1s19

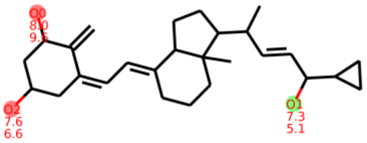

1tow

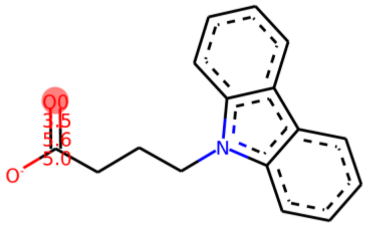

1tt1

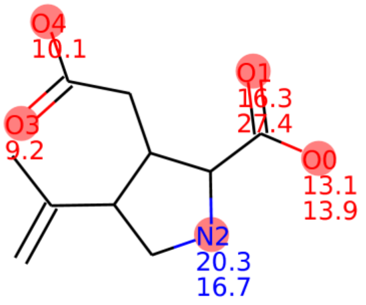


1u4d

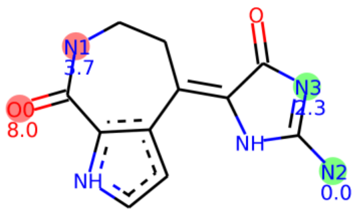


1ukz

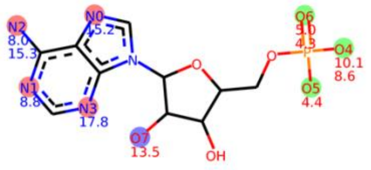


1ulb

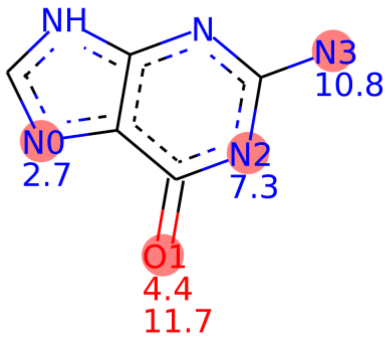


1unl

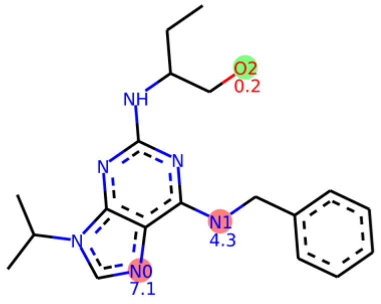


1uou

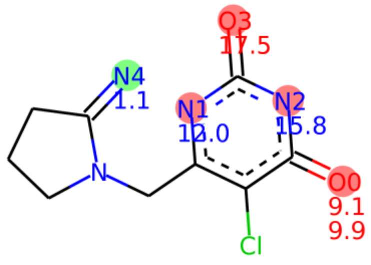


1v0p

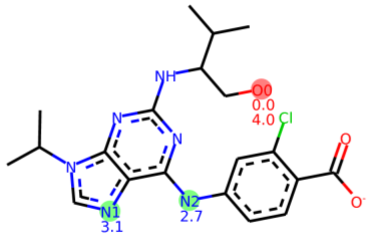


1w1p

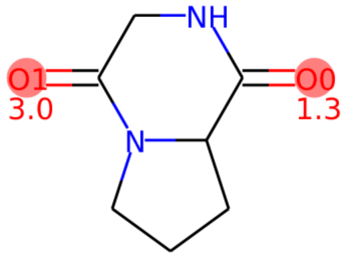


1w2g

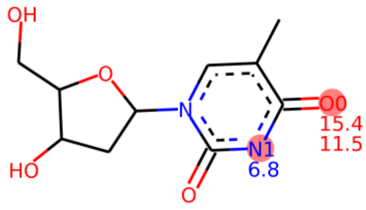


1x8x

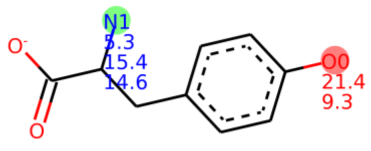


1ydr

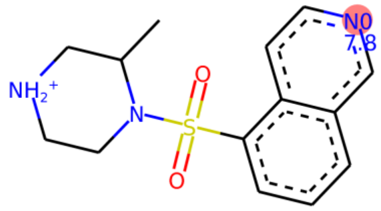


1yds

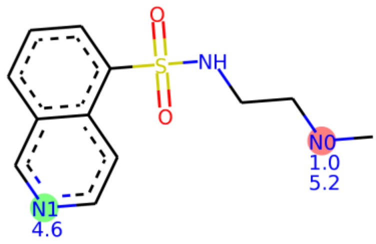

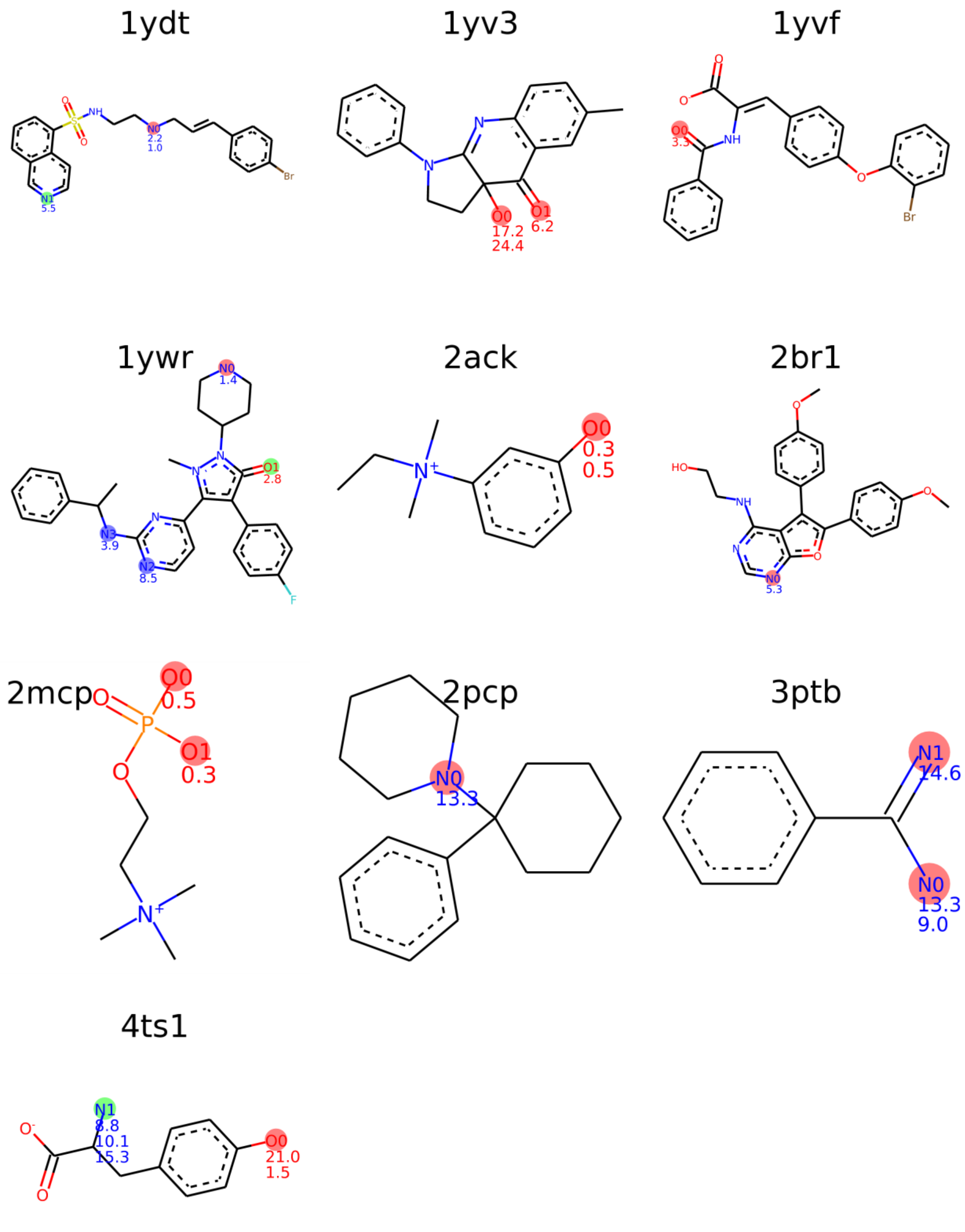
1ydt
1yv3
1yvf
1ywr
2ack
2br1
2mcp
2pcp
3ptb
4ts1

***Supplementary Table 6*** *Simulated structures classified by the number of structural anchors.*

| Number of structural anchors | PDB codes |
|---|---|
| 0 | 1c1b, 1f0s, 1g9v, 1hgh-2, 1jla, 1w1p, 1yvf, 2ack, 2mcp |
| 1 | 1a28, 1ai5, 1b9v, 1br6, 1cvu, 1dds, 1ezq, 1f0t, 1f0u, 1fcz, 1fm6, 1fvt, 1gm8, 1h1p, 1h1s, 1hgj, 1hwi, 1ivb, 1ivd, 1ive, 1ke5, 1l2s, 1lpz, 1lqd, 1ml1, 1mq6, 1mts, 1n2j, 1n2v, 1owe, 1pmn, 1q41, 1qhi, 1rob, 1tow, 1tt1, 1u4d, 1ulb, 1unl, 1uou, 1v0p, 1w2g, 1ydr, 1ydt, 1yv3, 1ywr, 2br1, 2pcp, 3ptb |
| 2 | 1exa, 1fcx, 1fh8, 1fh9, 1fhd, 1hgh-1, 1hgi, 1ivf, 1k1j, 1k3u, 1n46, 1of1, 1oyt, 1q1g, 1s19, 1x8x, 1yds, 4ts1 |
| 3 | 1l7f, 1m2z, 1ukz |

***Supplementary Table 7*** *Distribution of carbohydrate binding proteins based on the number of anchoring points.*

| No. anchoring points | No. complexes | % |
|---|---|---|
| 0 | 1 | 7.1 |
| 1 | 6 | 42.9 |
| 2 | 6 | 42.9 |
| 3 | 1 | 7.1 |

***Supplementary Figure 6*** *Standard deviation of $W_{QB}$ values within each cluster of hydrogen bonds selected based on Supplementary Figure 5 (marked in blue). Standard deviation of collected individual values separated by the number of hydrogen bonds in the group (marked in red).*

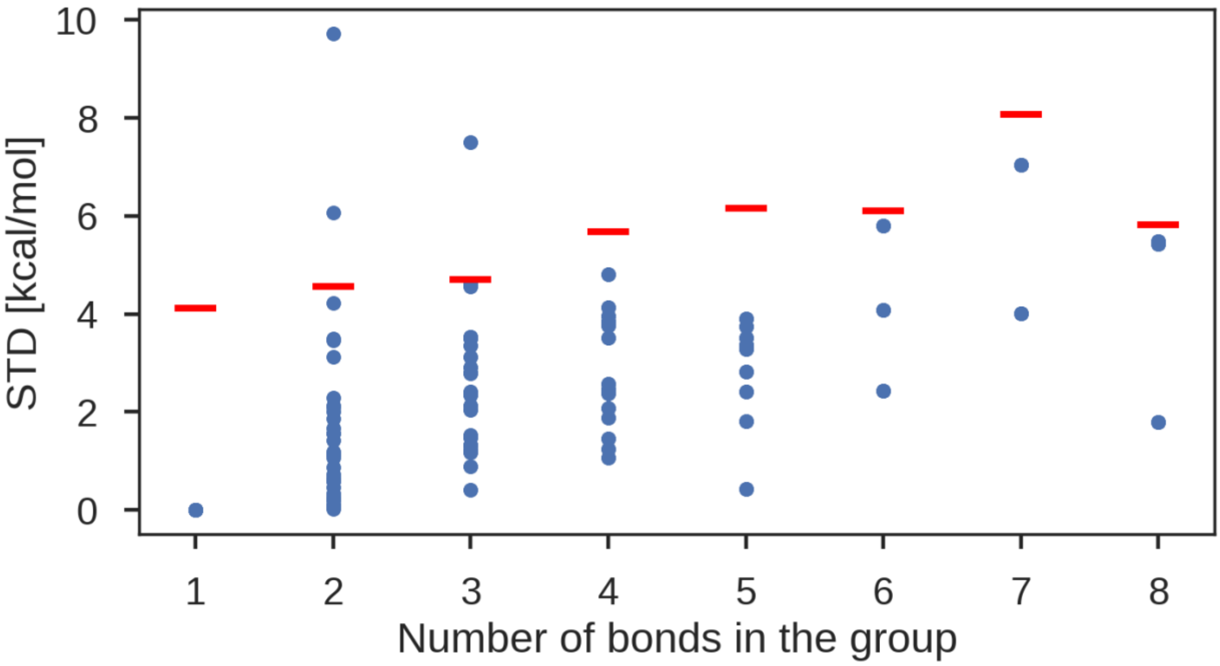


***Supplementary Figure 7*** *Depicted structures of ligand with PDB code 1n46 in the correct protonation state (left figure) in protonated form (middle figure). Atoms making hydrogen bonds are highlighted and the value listed below the atom correspond to $W_{QB}$ value of a bond made by this atom. Miscalculated protonation state not only is responsible for not forming an important interaction ($W_{QB}$=21.6 kcal/mol) but also significantly decreases the strength of 3 HBs in the local environment (from $W_{QB}$=19.9, 14.8 and 12.0 kcal/mol to $W_{QB}$=11.8, 11.9 and 9.4 kcal/mol, respectively). Both protonation states are predicted to exist at pH 7 (www.chemicalize.org). The right figure shows hydrogen bond network of considered scaffold.*

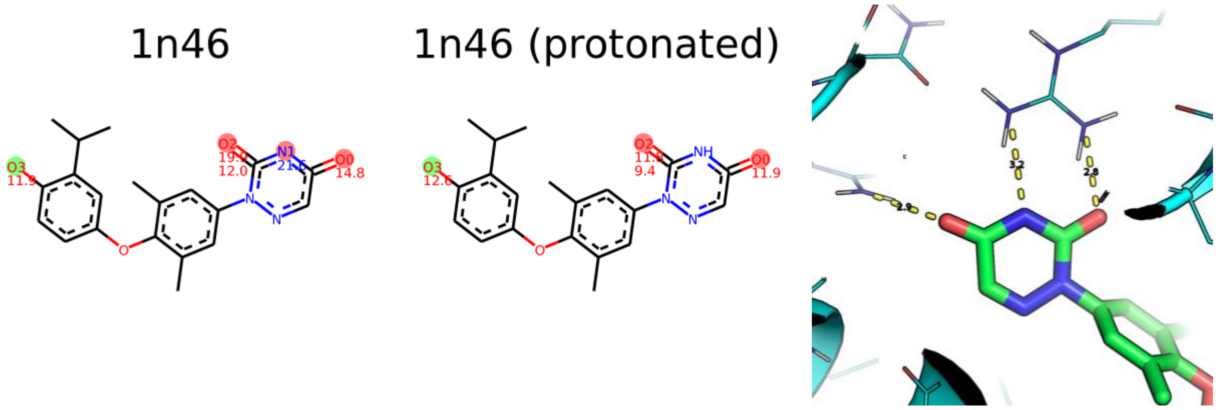

***Supplementary Figure 8** Binding free energy versus maximal $W_{QB}$ value for complexes with publicly available activity data. Binding energy was calculated based on: ΔG (blue), IC50 (green), $K_D$ (red) or $K_i$ (purple) measurements.*

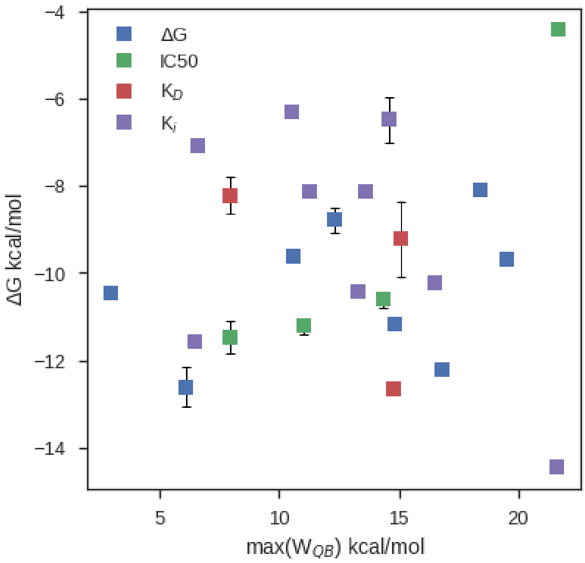


***Supplementary Figure 9** $W_{QB}$ value versus Solvent Accessible Surface Area (SASA) of the atom in the protein that makes the main hydrogen bond with ligand. All DUck-simulation data for Iridium DS (345 data points) are plotted.*

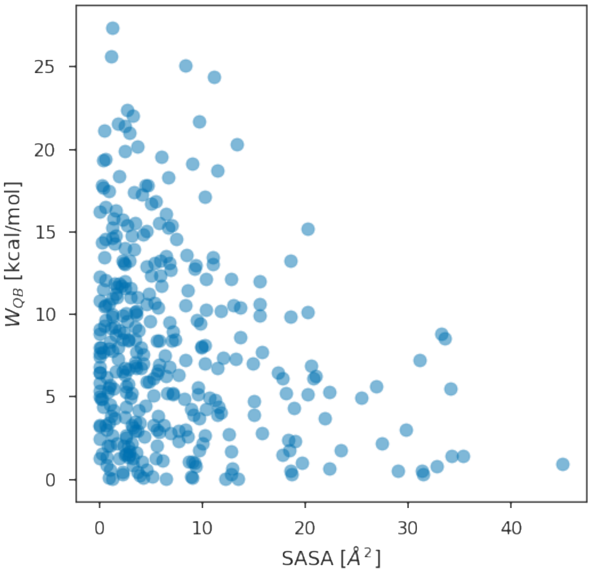


***Supplementary Table 8** Classification of complexes from SERAPhiC data set*

| | |
|---|---|
| Simulated structures | 1e2i_a, 1e2i_b, 1f5f, 1f8e, 1h46, 1k0e, 1mlw, 1ofz_a, 1ofz_b, 1r5y, 1sd1, 1tku, 1w1a, 1ynh, 2bkx, 2brt, 2f6x, 2fgq, 2hdq_a, 2hdq_b, 2i5x, 2iba, 2j5s, 2p1o, 2q6m, 2uy5, 3eko |
| No HB | 1sqn, 1ui0, 1uwc, 2cix, 2qwx, 3c0z, 3dsx |
| Metal ion | 1m2x, 1m3u, 1s5n, 1t0l, 1wog, 1x07, 1xfg, 1y2k, 1yv5, 2aie, 2fdv, 2ff2, 2gg7, 2gvv, 2rdr, 2v77, 2zvj |
| Additional ligand | 1fsg, 1pwm, 1yki, 2b0m, 2bl9 |

***Supplementary Figure 10*** *Depicted ligands from SERAPhiC DS, with corresponding PDB codes. Atoms that make hydrogen bonds with the protein were highlighted and the $W_{QB}$ value associated with that bond is written below the atom (multiple values in cases when the atom is making more than one hydrogen bond).*

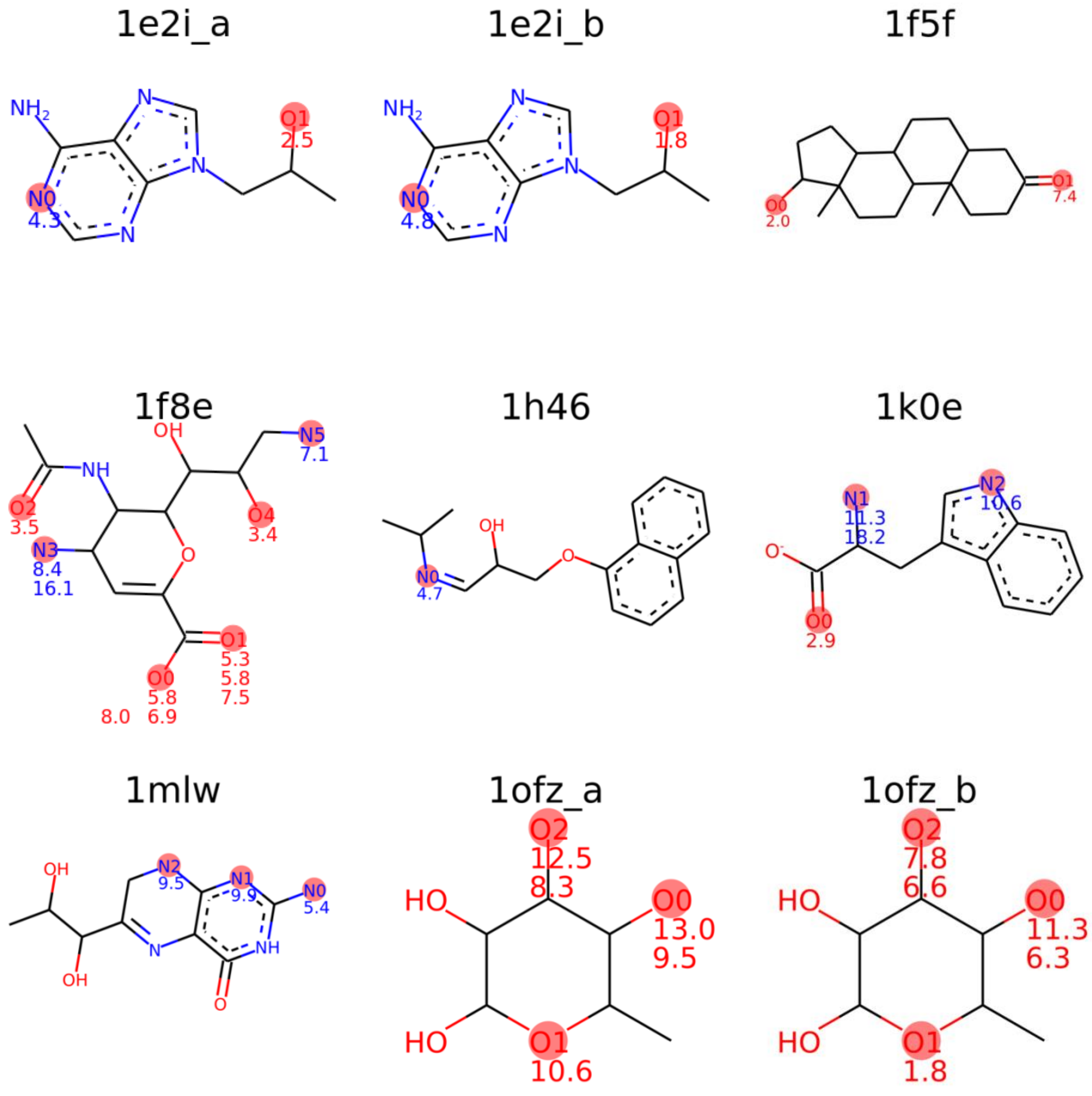

1r5y

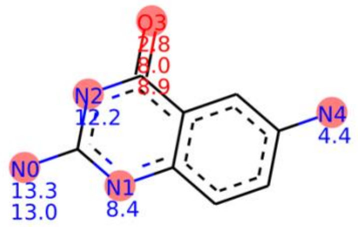

1sd1

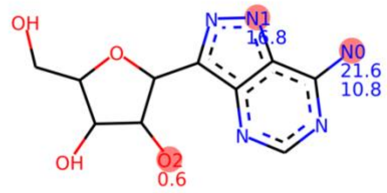

1tku

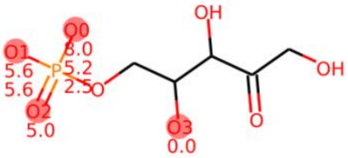

1w1a

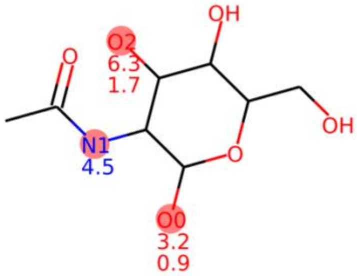

1ynh

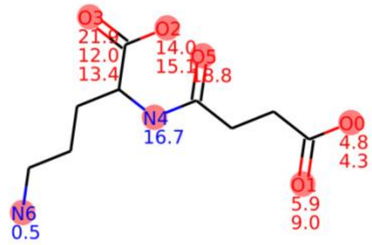

2bkx

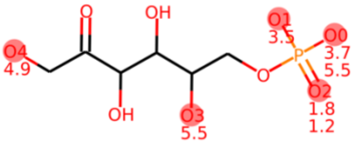

2brt

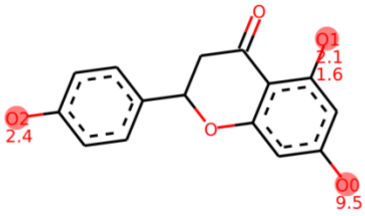

2f6x

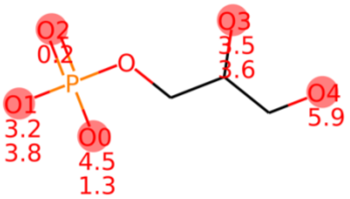

2fgq

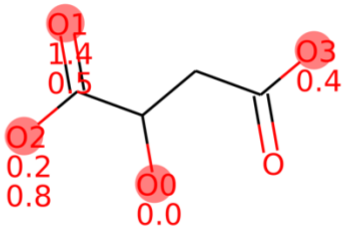

2hdq_a

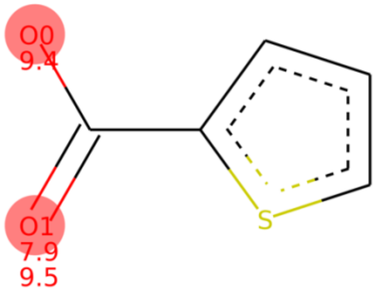

2hdq_b

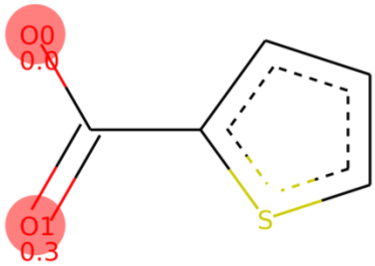

2i5x

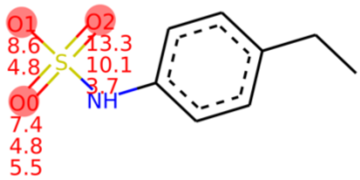

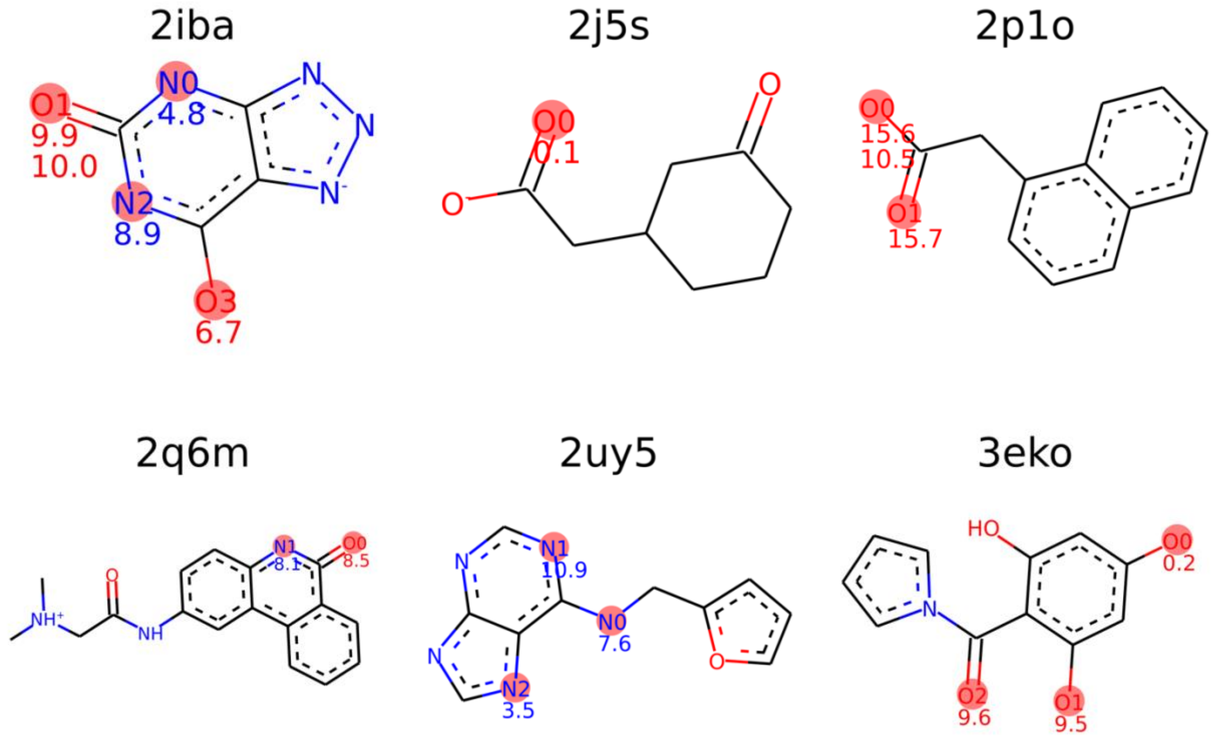


***Supplementary Table 9*** *List of hydrogen bonds for simulated systems from SERAPhiC DS. The bonds are classified by the complex's PDB code, residue number and atom name of protein's atom that makes the hydrogen bond. Table additionally contains, stability evaluation in a form of calculated $W_{QB}$ value, Mean_$W_{QB}$ value with standard deviation, the orientation of hydrogen bond (donor/acceptor function of protein's atom), and the bond character (information if the bond makes a salt bridge or neutral interaction).*

| PDB | RESIDUE | PROT_ATOM | # SMD | $W_{QB}$ [KCAL/MOL] | MEAN_WQB [KCAL/MOL] | STD [KCAL/MOL] | PROT (D/A) | SALT BRIDGE | NEUTRAL HB |
|---|---|---|---|---|---|---|---|---|---|
| **1E2I_A** | GLN125 | NE2 | 40 | 4.30 | 5.06 | 0.54 | D | 0 | 1 |
| | GLU83 | OE1 | 40 | 2.46 | 3.47 | 0.59 | A | 0 | 0 |
| **1E2I_B** | GLN125 | NE2 | 40 | 4.76 | 5.88 | 0.80 | D | 0 | 1 |
| | GLU83 | OE1 | 40 | 1.84 | 2.05 | 0.26 | A | 0 | 0 |
| **1F5F** | ASN82 | ND2 | 40 | 2.01 | 4.41 | 1.81 | D | 0 | 1 |
| | SER42 | OG | 40 | 7.36 | 7.83 | 0.29 | D | 0 | 1 |
| **1F8E** | ARG118 | NH1 | 40 | 5.78 | 5.93 | 0.12 | D | 1 | 0 |
| | ARG118 | NH2 | 40 | 6.91 | 7.13 | 0.17 | D | 1 | 0 |
| | ARG152 | NH2 | 40 | 3.52 | 4.49 | 0.65 | D | 0 | 0 |
| | ARG292 | NH1 | 40 | 5.34 | 5.91 | 0.53 | D | 1 | 0 |
| | ARG292 | NH2 | 40 | 5.76 | 6.93 | 0.68 | D | 1 | 0 |
| | ARG371 | NH1 | 40 | 7.51 | 8.57 | 0.69 | D | 1 | 0 |
| | ARG371 | NH2 | 40 | 7.97 | 9.11 | 0.75 | D | 1 | 0 |
| | ASP151 | OD1 | 40 | 8.35 | 9.60 | 0.77 | A | 0 | 0 |
| | GLU119 | OE2 | 40 | 16.14 | 16.37 | 0.29 | A | 0 | 0 |
| | GLU276 | OE1 | 40 | 3.35 | 3.52 | 0.18 | A | 0 | 0 |

| | | | | | | | | | |
|---|---|---|---|---|---|---|---|---|---|
| | GLU276 | OE2 | 40 | 7.09 | 7.91 | 0.63 | A | 1 | 0 |
| **1H46** | GLH207 | OE2 | 40 | 4.66 | 5.18 | 0.43 | D | 0 | 1 |
| **1K0E** | PHE46 | N | 40 | 2.91 | 4.06 | 0.67 | D | 0 | 0 |
| | SER242 | O | 40 | 10.58 | 13.85 | 1.91 | A | 0 | 1 |
| | SER36 | OG | 40 | 11.32 | 12.31 | 0.65 | A | 0 | 0 |
| | TYR43 | O | 40 | 18.18 | 19.61 | 0.84 | A | 0 | 0 |
| **1MLW** | GLY234 | O | 40 | 5.41 | 5.69 | 0.27 | A | 0 | 1 |
| | LEU236 | N | 40 | 9.88 | 11.07 | 1.17 | D | 0 | 1 |
| | LEU236 | O | 40 | 9.49 | 10.21 | 0.61 | A | 0 | 1 |
| **1OFZ_A** | ARG24 | NE | 40 | 13.04 | 13.75 | 0.54 | D | 0 | 0 |
| | ARG24 | NH2 | 40 | 10.63 | 10.81 | 0.16 | D | 0 | 0 |
| | GLU36 | OE1 | 40 | 12.52 | 12.77 | 0.15 | A | 0 | 0 |
| | GLU36 | OE2 | 40 | 9.50 | 9.84 | 0.27 | A | 0 | 0 |
| | TRP97 | NE1 | 40 | 8.27 | 8.62 | 0.20 | D | 0 | 1 |
| **1OFZ_B** | ARG226 | NE | 40 | 11.35 | 11.96 | 0.44 | D | 0 | 0 |
| | ARG226 | NH2 | 40 | 1.76 | 2.13 | 0.41 | D | 0 | 0 |
| | GLU238 | OE1 | 40 | 7.77 | 8.52 | 0.48 | A | 0 | 0 |
| | GLU238 | OE2 | 40 | 6.27 | 6.63 | 0.23 | A | 0 | 0 |
| | TRP298 | NE1 | 40 | 6.58 | 6.73 | 0.12 | D | 0 | 1 |
| **1R5Y** | ASH102 | OD1 | 40 | 13.25 | 14.04 | 0.52 | A | 0 | 1 |
| | ASH102 | OD2 | 40 | 8.38 | 9.14 | 0.46 | D | 0 | 1 |
| | ASP156 | OD1 | 40 | 12.95 | 16.89 | 2.34 | A | 0 | 0 |
| | ASP156 | OD2 | 40 | 12.21 | 12.91 | 0.42 | A | 0 | 0 |
| | CYS158 | SG | 40 | 2.80 | 3.21 | 0.39 | D | 0 | 1 |
| | GLN203 | NE2 | 40 | 7.98 | 10.16 | 1.72 | D | 0 | 1 |
| | GLY230 | N | 40 | 8.87 | 11.09 | 1.98 | D | 0 | 1 |
| | LEU231 | O | 40 | 4.36 | 4.77 | 0.29 | A | 0 | 1 |
| **1SD1** | ASP220 | OD1 | 40 | 21.57 | 22.13 | 0.55 | A | 0 | 0 |
| | ASP220 | OD2 | 40 | 16.80 | 17.12 | 0.31 | A | 0 | 0 |
| | ASP222 | OD1 | 40 | 10.84 | 11.16 | 0.28 | A | 0 | 0 |
| | MET196 | N | 40 | 0.59 | 1.78 | 1.08 | D | 0 | 1 |
| **1TKU** | ARG142 | NH1 | 40 | 8.03 | 9.14 | 0.99 | D | 1 | 0 |
| | ARG142 | NH2 | 40 | 5.59 | 7.06 | 1.14 | D | 1 | 0 |
| | CYS59 | SG | 4 | 0.00 | 0.09 | 0.10 | A | 0 | 1 |
| | HIE145 | N | 40 | 5.02 | 5.66 | 0.58 | D | 0 | 0 |
| | THR146 | N | 40 | 5.25 | 6.03 | 0.49 | D | 0 | 0 |
| | THR146 | OG1 | 40 | 2.53 | 4.52 | 1.20 | D | 0 | 0 |
| | THR85 | OG1 | 40 | 5.61 | 7.38 | 1.23 | D | 0 | 0 |
| **1W1A** | ARG166 | N | 40 | 3.19 | 3.79 | 0.42 | D | 0 | 0 |
| | ASP73 | OD2 | 40 | 6.27 | 7.06 | 0.83 | A | 0 | 0 |
| | HID124 | NE2 | 40 | 4.46 | 5.48 | 0.86 | A | 0 | 1 |
| | HIE128 | NE2 | 40 | 0.93 | 2.68 | 1.08 | D | 0 | 1 |
| | HIP222 | NE2 | 40 | 1.65 | 1.97 | 0.21 | D | 0 | 0 |

| | | | | | | | | | |
|---|---|---|---|---|---|---|---|---|---|
| **1YNH** | ALA19 | N | 40 | 4.81 | 5.81 | 0.94 | D | 0 | 0 |
| | ARG138 | NH1 | 40 | 13.99 | 14.25 | 0.18 | D | 1 | 0 |
| | ARG212 | NH1 | 40 | 15.07 | 18.31 | 2.19 | D | 1 | 0 |
| | ARG212 | NH2 | 40 | 21.89 | 25.36 | 2.03 | D | 1 | 0 |
| | ASN110 | OD1 | 40 | 0.52 | 0.65 | 0.15 | A | 0 | 0 |
| | ASN25 | ND2 | 40 | 12.02 | 12.25 | 0.18 | D | 0 | 0 |
| | ASN359 | ND2 | 40 | 4.34 | 4.37 | 0.03 | D | 0 | 0 |
| | ASN359 | O | 40 | 16.66 | 16.96 | 0.17 | A | 0 | 1 |
| | HIE137 | NE2 | 40 | 18.84 | 20.12 | 1.14 | D | 0 | 1 |
| | LEU21 | N | 40 | 5.92 | 6.31 | 0.47 | D | 0 | 0 |
| | SER22 | OG | 40 | 13.42 | 13.78 | 0.24 | D | 0 | 0 |
| | SER28 | OG | 40 | 9.03 | 9.61 | 0.39 | D | 0 | 0 |
| **2BKX** | ARG167 | NH2 | 40 | 3.70 | 5.38 | 1.41 | D | 1 | 0 |
| | GLY38 | N | 40 | 5.48 | 5.71 | 0.26 | D | 0 | 0 |
| | HID138 | NE2 | 40 | 5.52 | 6.33 | 0.52 | A | 0 | 1 |
| | LYS202 | NZ | 40 | 3.47 | 5.27 | 1.11 | D | 1 | 0 |
| | THR36 | N | 40 | 4.91 | 5.16 | 0.34 | D | 0 | 1 |
| | THR39 | N | 40 | 1.80 | 2.37 | 0.37 | D | 0 | 0 |
| | THR39 | OG1 | 40 | 1.16 | 1.67 | 0.73 | D | 0 | 0 |
| **2BRT** | GLU306 | OE2 | 40 | 9.50 | 11.30 | 1.73 | A | 0 | 0 |
| | SER236 | N | 40 | 2.09 | 2.56 | 0.45 | D | 0 | 1 |
| | TYR142 | OH | 40 | 2.40 | 4.62 | 1.77 | A/D | 0 | 1 |
| | VAL235 | N | 40 | 1.62 | 2.39 | 0.76 | D | 0 | 1 |
| **2F6X** | ASN1216 | N | 40 | 4.53 | 5.38 | 0.78 | D | 0 | 0 |
| | ASN1216 | ND2 | 40 | 1.33 | 3.62 | 1.40 | D | 0 | 0 |
| | GLU1167 | OE2 | 40 | 3.50 | 3.99 | 0.46 | A | 0 | 0 |
| | GLY1170 | N | 40 | 3.16 | 4.87 | 1.01 | D | 0 | 0 |
| | GLY1195 | N | 40 | 3.84 | 4.69 | 0.84 | D | 0 | 0 |
| | GLY1215 | N | 40 | 0.17 | 1.16 | 0.89 | D | 0 | 0 |
| | LYS1011 | NZ | 40 | 5.89 | 6.29 | 0.26 | D | 0 | 0 |
| | TYR1165 | OH | 40 | 3.56 | 4.18 | 0.65 | D | 0 | 1 |
| **2FGQ** | ARG133 | NH1 | 40 | 0.04 | 0.15 | 0.12 | D | 0 | 0 |
| | ARG133 | NH2 | 40 | 1.42 | 1.81 | 0.36 | D | 1 | 0 |
| | ARG75 | NH1 | 28 | 0.20 | 1.44 | 1.26 | D | 1 | 0 |
| | ARG75 | NH2 | 40 | 0.45 | 1.48 | 0.96 | D | 1 | 0 |
| | THR102 | OG1 | 40 | 0.45 | 0.99 | 0.36 | D | 0 | 0 |
| | THR36 | OG1 | 40 | 0.79 | 1.62 | 0.56 | D | 0 | 0 |
| **2HDQ_A** | ARG148 | NH1 | 40 | 9.39 | 10.35 | 0.66 | D | 1 | 0 |
| | ARG148 | NH2 | 40 | 7.94 | 9.42 | 0.86 | D | 1 | 0 |
| | LYS290 | NZ | 40 | 9.53 | 9.84 | 0.24 | D | 1 | 0 |
| **2HDQ_B** | GLY320 | N | 12 | 0.03 | 1.37 | 1.16 | D | 0 | 0 |
| | SER212 | N | 40 | 0.31 | 0.54 | 0.20 | D | 0 | 0 |
| **2I5X** | ALA1906 | N | 40 | 7.38 | 7.94 | 0.39 | D | 0 | 0 |

| | | | | | | | | | |
|---|---|---|---|---|---|---|---|---|---|
| | ARG1910 | N | 40 | 8.56 | 8.68 | 0.08 | D | 1 | 0 |
| | ARG1910 | NE | 40 | 4.81 | 5.13 | 0.21 | D | 1 | 0 |
| | ARG1910 | NH2 | 40 | 4.81 | 5.10 | 0.19 | D | 1 | 0 |
| | CYS1904 | SG | 40 | 13.28 | 13.89 | 0.58 | D | 0 | 0 |
| | GLY1909 | N | 40 | 10.11 | 10.23 | 0.07 | D | 0 | 0 |
| | SER1905 | N | 40 | 5.47 | 6.04 | 0.55 | D | 0 | 0 |
| | VAL1908 | N | 40 | 3.65 | 4.72 | 0.84 | D | 0 | 0 |
| **2IBA** | ARG176 | NH1 | 40 | 4.83 | 5.67 | 0.62 | D | 0 | 0 |
| | ARG176 | NH2 | 40 | 9.87 | 11.19 | 0.87 | D | 0 | 0 |
| | GLN228 | NE2 | 40 | 6.71 | 6.99 | 0.20 | D | 0 | 1 |
| | GLN228 | OE1 | 40 | 8.94 | 9.25 | 0.27 | A | 0 | 1 |
| | VAL227 | N | 40 | 9.97 | 10.57 | 0.56 | D | 0 | 1 |
| **2J5S** | HIP121 | NE2 | 40 | 0.08 | 2.73 | 1.67 | D | 1 | 0 |
| **2P1O** | ARG403 | NE | 40 | 15.57 | 16.40 | 0.83 | D | 1 | 0 |
| | ARG403 | NH2 | 40 | 15.73 | 15.82 | 0.07 | D | 1 | 0 |
| | SER438 | OG | 40 | 10.51 | 10.83 | 0.24 | D | 0 | 0 |
| **2Q6M** | GLY461 | N | 40 | 8.49 | 9.40 | 0.54 | D | 0 | 1 |
| | GLY461 | O | 40 | 8.07 | 8.61 | 0.49 | A | 0 | 1 |
| **2UY5** | TYR214 | OH | 40 | 7.55 | 9.39 | 1.25 | A | 0 | 1 |
| | ASP155 | OD2 | 40 | 10.95 | 11.62 | 0.61 | A | 0 | 0 |
| | TYR214 | OH | 40 | 3.52 | 4.56 | 0.62 | D | 0 | 1 |
| **3EKO** | ASN51 | ND2 | 40 | 0.24 | 1.30 | 0.93 | D | 0 | 1 |
| | ASP93 | OD2 | 40 | 9.51 | 12.36 | 1.67 | A | 0 | 0 |
| | THR184 | OG1 | 40 | 9.64 | 10.14 | 0.42 | D | 0 | 1 |

***Supplementary Figure 11** Mean-$W_{QB}$ value with standard deviation plotted against $W_{QB}$ value.*

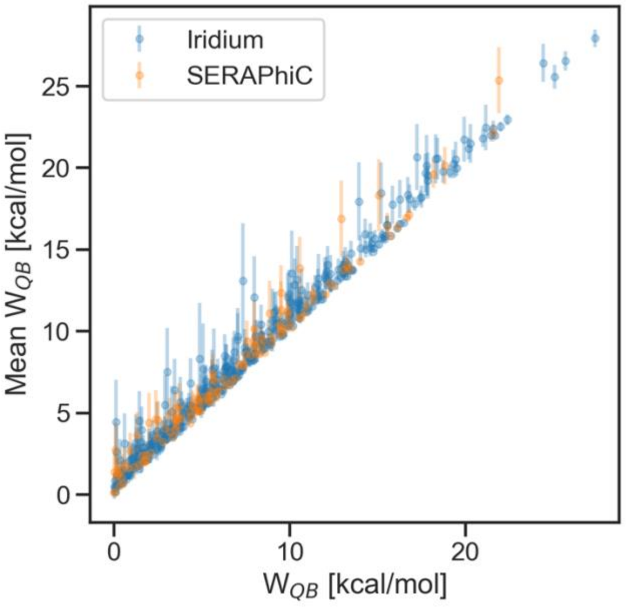
